# Beyond chromatin accessibility: bulk ATAC-seq as an integrative assay to portray genomes and epigenomes

**DOI:** 10.1101/2025.01.18.633115

**Authors:** Islem Toumi, Chaimae Kham, Lucille Stuani, Laurent Lecam, Pierre-François Roux

## Abstract

Assay for transposase-accessible chromatin using sequencing (ATAC-seq) is a cornerstone for epigenomic profiling, yet its potential for genomic characterization remains poorly explored. Here, we systematically benchmarked bulk ATAC-seq against whole-genome sequencing (WGS) to assess its capacity for detecting small variants, copy number variations (CNVs), telomere-associated repeat content, and mitochondrial single nucleotide polymorphisms in cancer cells. Using paired datasets from patient-derived melanoma cell lines and from TCGA primary brain tumors, we demonstrated that ATAC-seq achieves high precision in small variants detection within accessible regions supporting cohort-scale genotyping and genetic stratification, robustly resolves CNVs in the nuclear genome, and support high-coverage mitogenome profiling, with strong concordance to WGS at standard sequencing depths. Notably, we present the first systematic evaluation of telomere-associated repeat content by ATAC-seq, revealing its untapped potential for studying genome stability. By bridging genomic and epigenomic insights into a single genome-wide approach, bulk ATAC-seq emerges as a cost-effective and versatile tool poised to transform cancer research and to support integrative molecular profiling in clinical settings.

## INTRODUCTION

Chromatin accessibility is a central determinant of genome function, governing how regulatory information encoded in DNA is interpreted to establish cell identity, support developmental programs, and enable adaptive responses to environmental cues. Perturbation of chromatin accessibility landscapes is tightly linked to disease states, and has emerged as a defining feature of cancer, where epigenetic and transcriptional deregulation contribute to tumor initiation, progression, and therapy resistance(1, 2). The advent of ATAC-seq (Assay for Transposase-Accessible Chromatin using sequencing), which couples genome-wide Tn5 tagmentation to high-throughput sequencing, has transformed the study of chromatin organization by enabling rapid, high-resolution profiling of accessible chromatin from limited material. Since its introduction as a streamlined alternative to DNase-seq and FAIRE-seq, ATAC-seq has been extensively validated as a versatile method to map regulatory elements, infer transcription factor activity, and characterize enhancer landscapes across diverse biological systems(3). Continuous technical refinements have further broadened its applicability to heterogeneous and clinically derived samples, reinforcing its importance for translational research(4, 5).

Cancer, however, is not solely an epigenetic disease. Tumor development arises from the combined action of genomic and epigenomic alterations that together shape cellular fitness, plasticity, and evolutionary trajectories. Somatic point mutations, copy number variations (CNVs), structural rearrangements, and chromosomal instability constitute core hallmarks of cancer genomes and act in concert to drive malignant behavior. In the clinical setting, systematic interrogation of these genomic alterations is indispensable for diagnosis, classification, and treatment selection, particularly within precision oncology frameworks. Mutations affecting oncogenes such as *BRAF, EGFR*, or *MYC* directly inform targeted therapeutic strategies and have substantially improved patient outcomes (6, 7). Accordingly, tumor genomic profiling currently relies on a combination of resource-intensive technologies - including comparative genomic hybridization, karyotyping, whole-genome and whole-exome sequencing, and fluorescence in situ hybridization - to capture complementary aspects of genome structure and variation(8). While these approaches provide detailed access to the tumor genome, they offer limited insight into the epigenomic layer that regulates gene expression and cellular state. Yet, epigenomic dysregulation is now recognized as a fundamental driver of cancer therapeutic response and relapse (9). Altered chromatin accessibility, histone modifications, and DNA methylation patterns, indeed, rewire regulatory circuits, enabling oncogene activation, tumor suppressor silencing, and transcriptional plasticity. In particular, aberrant chromatin accessibility can directly promote oncogenic programs through enhancer hijacking, super-enhancer reorganization, and transcription factor rewiring (10–12). Beyond tumor initiation, epigenomic states critically influence tumor evolution under therapeutic pressure. Chromatin accessibility is closely tied to drug sensitivity and resistance, and epigenomic features can serve as predictive biomarkers of clinical outcome(13, 14). Consistently, therapy-resistant cells often undergo epigenetic reprogramming, entering drug-tolerant states that ultimately seed relapse. Despite this central role in cancer biology and treatment response, epigenomic profiling remains largely absent from routine clinical workflows, in part because accessible and cost-effective technologies capable of integrating genomic and epigenomic information are still lacking.

Identifying a single experimental approach capable of capturing both genome structure and regulatory state would substantially streamline molecular profiling of cancer samples and fill a major gap in current cancer diagnostics. Because of its intrinsic properties, ATAC-seq, occupies a unique position in this landscape. Although primarily developed as an epigenomic assay, ATAC-seq generates genome-wide sequencing data that inherently retain information about underlying DNA sequence. This raises the possibility that bulk ATAC-seq could serve as a dual-purpose platform for joint genomic and epigenomic characterization - provided that its genome-derived signals can be robustly interpreted and validated.

Beyond its canonical use for identifying accessible chromatin regions, ATAC-seq provides multiple complementary readouts of chromatin-encoded regulatory activity. Digital footprinting approaches leverage local patterns of chromatin accessibility to infer transcription factor binding at nucleotide resolution(15–18), providing direct insights into regulatory mechanisms controlling gene expression. Additionally, ATAC-seq fragment length distributions report nucleosome occupancy, enabling inference of nucleosome positioning and higher-order chromatin organization (19, 20). Extending ATAC-seq to single cells has shifted the field from population-averaged accessibility profiles to direct, cell-by-cell interrogation of chromatin regulation, enabling analysis of regulatory heterogeneity within complex tissues and tumor microenvironments (21). This capability is particularly relevant in cancer, where malignant cells and surrounding stromal and immune populations occupy distinct - and often rapidly evolving - epigenomic states. By capturing accessibility and transcription factor footprinting signatures at single-cell resolution, scATAC-seq reveals regulatory landscapes with a granularity that is inaccessible in bulk assays(22). Together with multi-modal and integrative strategies, these developments have broadened the scope of precision oncology(23). scATAC-seq has proven particularly powerful for identifying rare cell states, discovering biomarkers, and guiding personalized therapeutic strategies(24, 25). More recently, spatial ATAC-seq approaches have enabled chromatin accessibility profiling in situ, uncovering spatially resolved regulatory programs in both healthy and diseased tissues(26).

Despite this breadth of information, applications of bulk ATAC-seq remain largely confined to epigenomic analyses such as accessible region annotation and regulatory element mapping (27). Yet bulk ATAC-seq datasets also contain substantial genome-derived information that has been comparatively underexploited. Several studies have demonstrated that ATAC-seq can support the detection of single nucleotide polymorphisms (SNPs) and short insertions or deletions (indels) with reasonable accuracy(28), enabling allele-specific analyses and chromatin accessibility quantitative trait mapping(29). In addition, the genome-wide coverage provided by ATAC-seq - even if uneven - can be leveraged to infer large-scale copy number variations (CNVs), particularly in cancer samples where CNVs are pervasive(30). While such genome-oriented applications have been explored in single-cell ATAC-seq, their robustness, scalability, and practical limits in bulk ATAC-seq remain insufficiently characterized(12, 30, 31). Similarly, because ATAC-seq captures mitochondrial DNA alongside nuclear DNA, it offers an opportunity to profile mitochondrial variation. Although single-cell ATAC-seq has been successfully used for this purpose(32, 33) systematic validation of mitochondrial analyses from bulk ATAC-seq remains largely absent.

One particularly overlooked application is the quantification of telomere-associated repeat content from bulk ATAC-seq. Telomeres - repetitive DNA tracts capping chromosome ends - are central to genome stability, replicative capacity, and cellular lifespan, and their dysfunction is a defining feature of cancer. Progressive telomere erosion imposes a proliferative barrier through replicative senescence, whereas many tumors circumvent this barrier by reactivating telomere maintenance programs, most commonly via telomerase (encoded by *TERT*) reactivation (34). Consequently, telomeric repeat abundance and composition are routinely assessed in cancer research using whole-genome sequencing or dedicated telomere assays, including qPCR-based approaches, telomere restriction fragment analysis, and fluorescence in situ hybridization-based methods. Because ATAC-seq inevitably captures fragments encompassing telomeric repeat motifs, these reads have been proposed as a potential surrogate for telomere length measurements. However, recent work has demonstrated that telomere-motif reads in ATAC-seq are intrinsically ambiguous (35): they can originate not only from telomeres, but also from subtelomeric domains, interstitial telomeric sequences, and telomere-adjacent repeat tracts enriched in telomeric variant repeats. As a result, telomere-associated signal in ATAC-seq may reflect broader chromatin-state or cell-cycle-dependent accessibility features rather than telomere length itself. To date, no systematic effort has quantified how telomeric repeat content inferred from bulk ATAC-seq relates to established WGS-based measurements after accounting for these confounding sources.

More generally, the extraction of genome-level information from bulk ATAC-seq is hindered by the lack of reference-grade benchmarks defining what can - and cannot - be reliably recovered under realistic sequencing depth and signal-to-noise regimes. Unlike WGS, ATAC-seq samples the genome unevenly, concentrating reads in accessible regions while leaving other loci sparsely covered. Consequently, while small-variant and CNV pipelines have been repurposed for ATAC-seq, their performance depends strongly on sequencing depth, local accessibility, and computational choices, and reported accuracies remain difficult to compare across studies. In particular, standardized evaluation frameworks are still missing for mitochondrial variation and telomere-associated repeat signal in bulk ATAC-seq, underscoring the need for systematic comparisons against matched gold-standard assays to define the practical operating range of bulk ATAC-seq as a dual-purpose genomic-epigenomic readout.

In this study, we leverage paired WGS and bulk ATAC-seq datasets to systematically benchmark the recovery of small variants, copy number alterations, telomere-associated repeat content, and mitochondrial variation from ATAC-seq. Our results demonstrate that bulk ATAC-seq is not only a powerful tool for epigenomic profiling but also a versatile platform for genomic characterization of tumor samples. By defining the strengths, limitations, and practical operating range of genome-derived signals in bulk ATAC-seq, we show that ATAC-seq can support robust epigenomic profiling while retaining informative readouts of genome variation and structure. Together, these findings position bulk ATAC-seq as a genuinely integrative assay that bridges genomic and epigenomic characterization, with direct relevance for cancer research and precision oncology.

## MATERIAL AND METHODS

### Data source and pre-processing

Matched raw bulk ATAC-seq, H3K27ac ChIP-seq, RNA-seq, and WGS data from ten patient-derived melanoma cell lines (A375, MM001, MM011, MM029, MM031, MM047, MM057, MM074, MM087, and MM099) were obtained from the European Genome-Phenome Archive (EGAS00001004136) and GEO (GSE134432, GSE142238, and GSE159965). All WGS libraries were paired-end 2 × 151 bp (mean 1.20 ± 0.72 × 10^9^ fragments; mean coverage 51 ± 31×), whereas ATAC-seq libraries were paired-end with heterogeneous read lengths (2 × 25 bp, 2 × 48 bp, or 2 × 50 bp; mean 90.8 ± 88.1 × 10^6^ reads; mean FRiP 41 ± 13%, Table 1). For downstream benchmarking analyses, we retained cell lines with WGS coverage > 30× and ATAC-seq datasets > 50 million fragments, resulting in a final set of six lines (A375, MM001, MM011, MM031, MM047, and MM074, Table 1). Following an initial quality assessment, WGS data were randomly downsampled with seqtk v1.3 (https://github.com/lh3/seqtk) to achieve 30× genome coverage. To minimize confounding effects of heterogeneous read lengths across datasets, ATAC-seq reads were hard-trimmed to 2 × 25 bp using Picard Tools v3.0.0 (http://broadinstitute.github.io/picard) prior to iterative downsampling with seqtk v1.3 to generate datasets with 10M, 20M, 30M, 40M, and 50M ATAC-seq fragments.

To extend the benchmarking beyond melanoma cell lines and evaluate performance in a less favorable and clinically realistic context, we additionally analyzed matched tumor WGS and bulk ATAC-seq data from The Cancer Genome Atlas glioblastoma (TCGA GBM) and low-grade glioma (TCGA LGG) cohorts (Table 1). This validation cohort comprised 22 primary tumors (13 LGG, 9 GBM), including *IDH1*^R132H^-mutant and *IDH*-wild-type cases. TCGA WGS tumor libraries were all paired-end 2 × 151 bp with a mean of 2.01 ± 0.15 × 10^9^ fragments and a mean coverage of 79 ± 7×, while TCGA ATAC-seq tumor libraries were all paired-end 2 × 76 bp with a mean of 156 ± 44 × 10^6^ fragments and a mean fraction of reads in peaks (FRiP) of 27 ± 12% (Table 1). WGS data were randomly down-sampled to achieve 30× genome coverage, while ATAC-seq data were iteratively down-sampled with seqtk v1.3 to generate datasets with 10M, 20M, 30M, 40M, and 50M fragments.

### ATAC-seq processing

To extract genetic features from bulk ATAC-seq, ATAC-seq data were processed with the standardized nf-core(36) sarek v3.4.0(37) pipeline using the GATK.GRCh38 reference genome. Briefly, input FASTQ files reads were preprocessed with fastp (38). Reads were then aligned using the BWA-MEM(39), coordinate-sorted with samtools(40), and processed following GATK Best Practices(41): PCR/optical duplicates were marked using GATK MarkDuplicates and base quality score recalibration was applied using GATK BaseRecalibrator/ApplyBQSR prior to variant discovery. Small variants (SNPs and short indels) were called in single-sample mode using DeepVariant(42), FreeBayes(43), GATK HaplotypeCaller(44), and bcftools/mpileup(45) with the default configurations provided by nf-core/sarek. The resulting VCFs were used for subsequent concordance analyses with 30× WGS-derived calls.

For epigenomic analyses, ATAC-seq data were processed using a standard epigenomic workflow consistent with ENCODE recommendations (46). Paired-end ATAC-seq reads were aligned with bowtie2 v2.5.1(47) (--very-sensitive-local) to the same GATK.GRCh38 reference genome. BAM files were coordinate-sorted and indexed, and alignments overlapping ENCODE blacklisted regions were removed using bedtools v2.30.0(48). Open chromatin peaks were identified using MACS3 v3.0.0(49) with default parameters. Super-enhancers were inferred using ROSE v1.3.2(11, 50) with a stitching distance of 12.5 kb (-s 12500) and a 2.5 kb exclusion window around transcription start sites (-t 2500). Transcription factor footprinting was performed using TOBIAS v0.16.1(51) together with the HOCOMOCO v11(52) motif database.

### WGS processing

WGS data were processed with the standardized nf-core(36) sarek v3.4.0(37) pipeline using the GATK.GRCh38 reference genome. Briefly, input FASTQ files reads were preprocessed with fastp(38). Reads were then aligned using the BWA-MEM, coordinate-sorted with samtools(40), and processed following GATK Best Practices(41): PCR/optical duplicates were marked using GATK MarkDuplicates and base quality score recalibration was applied using GATK BaseRecalibrator/ApplyBQSR prior to variant discovery. Small variants (SNPs and short indels) were called in single-sample mode using DeepVariant(42), FreeBayes(43), GATK HaplotypeCaller(44), and bcftools/mpileup(45) with the default configurations provided by nf-core/sarek. The resulting VCFs were used for subsequent concordance analyses with ATAC-seq-derived calls.

### ChIP-seq processing

H3K27ac ChIP-seq data from patient-derived melanoma cell-lines were processed using a standard epigenomic workflow consistent with ENCODE recommendations (46) for histone-mark ChIP-seq. Paired-end reads were aligned to the GATK.GRCh38 reference genome using bowtie2 v2.5.1 (47) with the --very-sensitive-local preset. Following alignment, BAM files were coordinate-sorted and indexed, and reads mapping to ENCODE blacklisted regions were removed using bedtools v2.30.0(48). Sharp H3K27ac enrichment regions were identified using MACS3 v3.0.0 (49) peak calling with default parameters. Super-enhancers were subsequently inferred from H3K27ac signal using ROSE v1.3.2 (11, 50), applying a stitching distance of 12.5 kb (-s 12500) and a 2.5 kb exclusion window around transcription start sites (-t 2500), as previously described.

### RNA-seq processing

RNA-seq data from patient-derived melanoma cell-lines were processed using the standardized nf-core (36) rnaseq v3.11.0 pipeline with the GATK.GRCh38 reference. Briefly, reads were preprocessed for adapter and quality trimming with TrimGalore (https://github.com/FelixKrueger/TrimGalore), and transcript quantification was performed using Salmon (53) in quasi-mapping mode against the corresponding GRCh38 transcriptome reference. For downstream analyses, we used the gene-level count matrix derived from Salmon quantification.

### Cancer-driving mutational signatures

To summarize cancer-relevant driver alterations, we generated Mutation Annotation Format (MAF) files from 30× WGS-derived small-variant callsets and performed cohort-specific oncoplot analyses. For the six melanoma cell lines retained for benchmarking, we used the Variant Effect Predictor (VEP)-annotated and filtered bcftools/mpileup VCF outputs and converted them to MAF format using vcf2maf v1.6.22, relying on Ensembl-VEP v111.0(54). Conversion was performed on the GRCh38 build. MAF files were then imported into R using maftools v2.24.0 (55). To prioritize biologically relevant variants, we applied a filtering strategy based on MAF annotations, retaining variants supported by literature and database evidence while excluding variants flagged as common. Filtered MAF objects were merged across samples and further restricted to COSMIC-tagged variants by selecting entries annotated with COSMIC(56) identifiers. Oncoplots were finally generated using a melanoma-focused driver gene panel (*TP53, BRAF, CDKN2A, NRAS, KIT, PTEN, CTNNB1*). The same VCF to MAF conversion and filtering workflow was applied to the TCGA LGG/GBM cohort, and oncoplots were generated using a glioblastoma- and low-grade glioma-relevant driver panel (*TP53, IDH1, IDH2, ATRX, PTEN, EGFR, NF1*).

#### *WGS, ATAC-seq, H3K27ac and RNA-seq* genome-wide signal concordance in patient-derived melanoma cell lines

To quantify concordance of genome-wide signal patterns across datasets in the patient-derived melanoma cell line cohort, we computed pairwise correlations between coverage tracks (log_10_(RPM)) generated with deepTools v3.5.2(57) bamCoverage, after binning the genome into fixed-width windows. GRCh38 autosomes were tiled into consecutive 1 kb bins. For each normalized signal track, signal was extracted over all bins using genomation v1.40.1(58) R package. Track-to-track similarity was assessed using Pearson correlation computed across all bins. Correlation matrices were visualized as clustered heatmaps, using correlation distance for rows and columns and Ward D^2^ hierarchical clustering.

#### *Integrative analysis based on Partial Least Square* in patient-derived melanoma cell lines

A block partial least squares (PLS) analysis was performed using the mixOmics v6.28.0(59) R package to integrate and explore relationships between multiple datasets capturing distinct molecular layers of the patient-derived melanoma cell lines. Chromatin accessibility (ATAC-seq) was represented by two blocks summarizing signals over super-enhancers and enhancers. Likewise, H3K27ac ChIP-seq profiles were encoded as two blocks summarizing signal over super-enhancers and enhancers. Transcription factor binding predictions were included as a dedicated block using scaled log_10_-transformed counts of predicted binding events to provide a functional proxy for regulatory activity. Gene expression was included as an additional block using normalized RNA-seq expression values. To relate these regulatory layers to genomic alterations, we also considered a block summarizing oncogenic driver events derived from 30× WGS. In the unsupervised block PLS integration performed in regression mode, this WGS driver block was treated as an illustrative outcome. The block PLS model was used to estimate latent components capturing the covariance structure shared across blocks, enabling the identification of coordinated patterns linking chromatin accessibility, H3K27ac activity, transcriptional programs, and the oncogenic driver landscape.

#### Small variants comparison

To evaluate the consistency and overlap of small variants (SNPs and indels) detected by ATAC-seq and 30× WGS, VCF files generated from both assays were analyzed using the MutationalPatterns v3.14.0(60) R package. Variant calls obtained from ATAC-seq were compared to 30× WGS-derived calls, treated as the gold-standard reference set. Overlapping variants were defined as calls matching genomic coordinates and alleles between datasets, and overlaps were visualized using proportional Venn diagrams drawn with gVenn (https://github.com/ckntav/gVenn). For readability, proportional Venn diagrams were generated as sample-wise, pairwise comparisons between 30× WGS and a given ATAC-seq sequencing depth, displaying only the variants detected in WGS. To quantify the accuracy of ATAC-seq variant calling relative to WGS, precision, recall, and F1 scores were computed separately for SNPs and indels using benchmarking summaries produced with hap.py v0.3.15 (https://github.com/illumina/hap.py). True positives (TP) were defined as ATAC-seq calls also detected in 30× WGS, false positives (FP) as ATAC-seq calls not detected in 30× WGS, and false negatives (FN) as 30× WGS calls not detected in ATAC-seq; precision, recall, and F1 score were computed from TP, FP, and FN as follow:

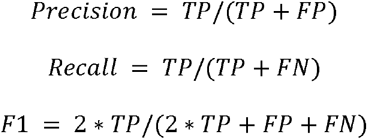

#### CNV calling and comparison

Copy-number profiling from bulk ATAC-seq and matched 30× WGS was performed in R v4.4.0 using QDNAseq v1.40.0(61) with GRCh38 annotation at 100 kb resolution. For each sample, CNV inference was applied to each downsampled ATAC-seq BAMs corresponding to 10M-50M fragments and to the match 30× WGS BAM. Read counts were computed with binReadCounts in paired-end mode. To ensure identical binning across assays while preserving assay-specific signal distributions, we first constructed two separate multi-sample QDNAseq objects: one containing all 30× WGS samples and one containing all ATAC-seq samples The QDNAseq preprocessing pipeline consisted of: (i) applyFilters with both residual and blacklist filtering enabled; (ii) estimation of technical biases with estimateCorrection followed by GC/mappability correction using correctBins; (iii) library-size normalization with normalizeBins; and (iv) outlier smoothing using smoothOutlierBins. Segmentation was performed on variance-stabilized signal using segmentBins with an Anscombe transform and local smoothing over 5 consecutive bins, followed by segment-level normalization with normalizeSegmentedBins. Discrete CNV calls were produced using callBins with the cutoff method and symmetric log_2_ thresholds (loss = −0.4, gain = +0.4) to assign per-bin states. CNV calling was performed independently for 30× WGS and ATAC-seq, and the processed objects were then merged into a single combined object to enable direct bin-level comparisons across assays and ATAC-seq depths. To reduce technical artefacts and sparse-bin effects in the multi-sample object, bins with zero read counts in at least one ATAC-seq library were removed prior to CNV calling. After calling, bins with extreme or missing copy-number estimates were excluded from downstream comparisons. For visualization and concordance analyses, per-bin log_2_(CNV) values from ATAC-seq and 30× WGS were extracted for each sample and depth, merged by genomic bin, and displayed using hexagonal binning density plots. To quantify agreement between assays, Pearson correlation coefficients were computed between 30× WGS- and ATAC-seq-derived per-bin log_2_(CNV) values for each matched sample and ATAC-seq depth. To evaluate CNV calling performance using 30× WGS as ground truth, QDNAseq call states were compared directly between ATAC-seq and 30× WGS for matched samples at each depth. Calls were collapsed into three clinically interpretable categories: loss (<2N), neutral (2N), and gain (>2N). For each CNV category, 30× WGS calls were used as the ground truth and ATAC-seq calls as the prediction, and classification was performed at the bin level: true positives (TP) were bins assigned to the category in both 30× WGS and ATAC-seq, false negatives (FN) were bins assigned to the category in 30× WGS but not in ATAC-seq, false positives (FP) were bins assigned to the category in ATAC-seq but not in 30× WGS, and true negatives (TN) were bins not assigned to the category in either assay; from these counts, precision, recall and accuracy were computed for each sample and ATAC-seq depth as follow:

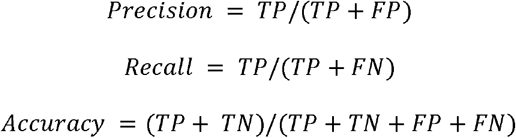

#### Telomere content computation and comparison

Telomere-associated repeat content was quantified from bulk ATAC-seq (10M-50M fragments) and matched 30× WGS using TelomereHunter v1.1.0(62), which outputs per-cytoband repeat spectrum tables and library read counts. For each sample, TelomereHunter spectra were parsed to extract counts for a panel of canonical and variant telomeric motifs and their reverse-complement (TTAGGG/CCCTAA, TGAGGG/CCCTCA, TCAGGG/CCCTGA, TTGGGG/CCCCAA). To enable cross-platform comparisons while accounting for library size differences, we generated library-size-scaled spectrum. To focus the analysis on telomeres and exclude intrachromosomal pseudo- and interstitial telomeric repeats, GRCh38 cytoband annotations were used. For each telomeric band, motif counts were summarized at the level of each motif. Bands with zero signal in either assay were excluded, and comparisons were performed on log-transformed values to stabilize variance. Concordance between assays was quantified using Pearson correlation between 30× WGS and ATAC-seq log-transformed motif abundance for each sample, and correlation coefficients and associated *p*-values were summarized as a function of sequencing depth and motif class. For the TCGA LGG/GBM cohort, we investigated whether telomere-associated repeat signal inferred from ATAC-seq was associated with *TERT* expression. *TERT* expression values were extracted from TCGA GDC portal(63). Canonical TTAGGG/CCCTAA motif counts were restricted to telomeres and summarized per sample and ATAC-seq depth. Associations with *TERT* expression were assessed using log-transformed telomere-associated signal and visualized across ATAC-seq depths. Samples were additionally stratified into *TERT* ON (expressed) versus *TERT* OFF (not expressed) groups as defined in (64).

#### Mitogenome variant calling, VAF profiling, and coverage analysis

Mitochondrial allele counts were obtained from bulk ATAC-seq (10M-50M fragments) and matched 30× WGS using mgatk v0.7.0(33) in bulk mode. To derive a consistent mtDNA variant catalogue across all datasets, variant calling was performed jointly across libraries, treating each downsampled ATAC-seq dataset and each 30× WGS dataset as an individual pseudo-sample within a single callset. This joint calling strategy increases sensitivity for low-frequency alleles and ensures that per-sample VAFs are computed over the same set of candidate sites, enabling direct cross-platform and cross-depth comparisons. Per-position alternate allele frequencies were reconstructed for each possible substitution by combining strand-specific counts, restricting the analysis to positions with an unambiguous non-unassigned reference base. To identify high-confidence mtDNA variants while accounting for coverage heterogeneity, we computed variant-level quality metrics across samples, including a strand concordance score defined as the Pearson correlation between forward and reverse alternate counts across non-zero observations and a variance-to-mean ratio (VMR) of allele frequency. As suggested in (33), variants were retained if they were confidently detected in at least one sample, showed strand concordance ≥ 0.65, and had log_10_(VMR) > −2. Variant allele frequency (VAF) was then computed per sample as VAF = alt /(ref + alt + ⍰) from raw nucleotide counts, with ⍰ = 1 × 10^−4^ to avoid division by zero, and per-variant coverage was defined as ref + alt. Sample-by-variant VAF matrices were used to compare mtDNA variant profiles across assays and ATAC-seq depths. VAF profiles were visualized as heatmaps with Ward’s D^2^ clustering and correlation-based distance after filtering out low-information entries by masking sites with VAF < 2% and coverage < 1000× and retaining variants with VAF standard deviation ≥ 0.05 across samples. Global similarity between mtDNA variant profiles was further assessed by principal component analysis (PCA) on the VAF matrix after logit transformation of allele frequencies. To compare mitochondrial coverage patterns between assays, per-base mtDNA coverage tracks were generated at 1 bp resolution with deepTools v3.5.2 (57) and imported into R. Coverage values were log_10_-transformed and summarized into fixed-width bins along the mitogenome (20 bp). For the patient-derived melanoma cell lines cohort including a limited sample number, coverage profiles were visualized on a per-sample basis to display individual 30× WGS and ATAC-seq depth-dependent patterns. In contrast, for the TCGA LGG/GBM cohort, coverage profiles were summarized to provide a cohort-level view: for each ATAC-seq depth, a meta-coverage profile was computed across tumors by taking the median coverage per bin, together with an interquartile range (25^th^-75^th^ percentile) ribbon to capture between-sample variability. 30× WGS meta-coverage was computed analogously. These profiles were displayed in polar coordinates to represent coverage along the circular mitogenome. An external gene annotation ring was drawn using Ensembl v111.0(54) mitochondrial gene coordinates.

## RESULTS

### ATAC-seq links oncogenic alterations to coherent regulatory programs in patient-derived melanoma cell lines

We leveraged multi-omics datasets from 10 patient-derived melanoma cell lines, including matched bulk ATAC-seq, H3K27ac ChIP-seq, RNA-seq, and WGS, to evaluate the extent to which bulk ATAC-seq can support both epigenomic profiling and genome-oriented analyses (Fig. 1A). This cohort provides a controlled experimental setting in which chromatin accessibility, transcriptional output, and genomic alterations can be directly compared within the same biological samples. We focused on short-variant detection, CNV inference, telomere length estimation, and profiling of the mitogenome, while quantifying the impact of sequencing depth through systematic downsampling. For benchmarking, we retained six cell lines with WGS > 30× and ATAC-seq > 50M fragments (A375, MM001, MM011, MM031, MM047, MM074; Fig. 1B). Within this subset, WGS coverage ranged from 33× to 54× (78-130M fragments), whereas ATAC-seq yielded 66M to 310M fragments and displayed a broad range of signal-to-noise ratios, with fraction of reads in peaks (FRiP) values spanning 19.6% to 59.5% (Fig. 1B, Table 1). To control for heterogeneous read lengths across libraries, all ATAC-seq datasets were hard-trimmed to 2 x 25 bp prior to iterative downsampling, from 10M to 50M fragments.

**Figure 1.**
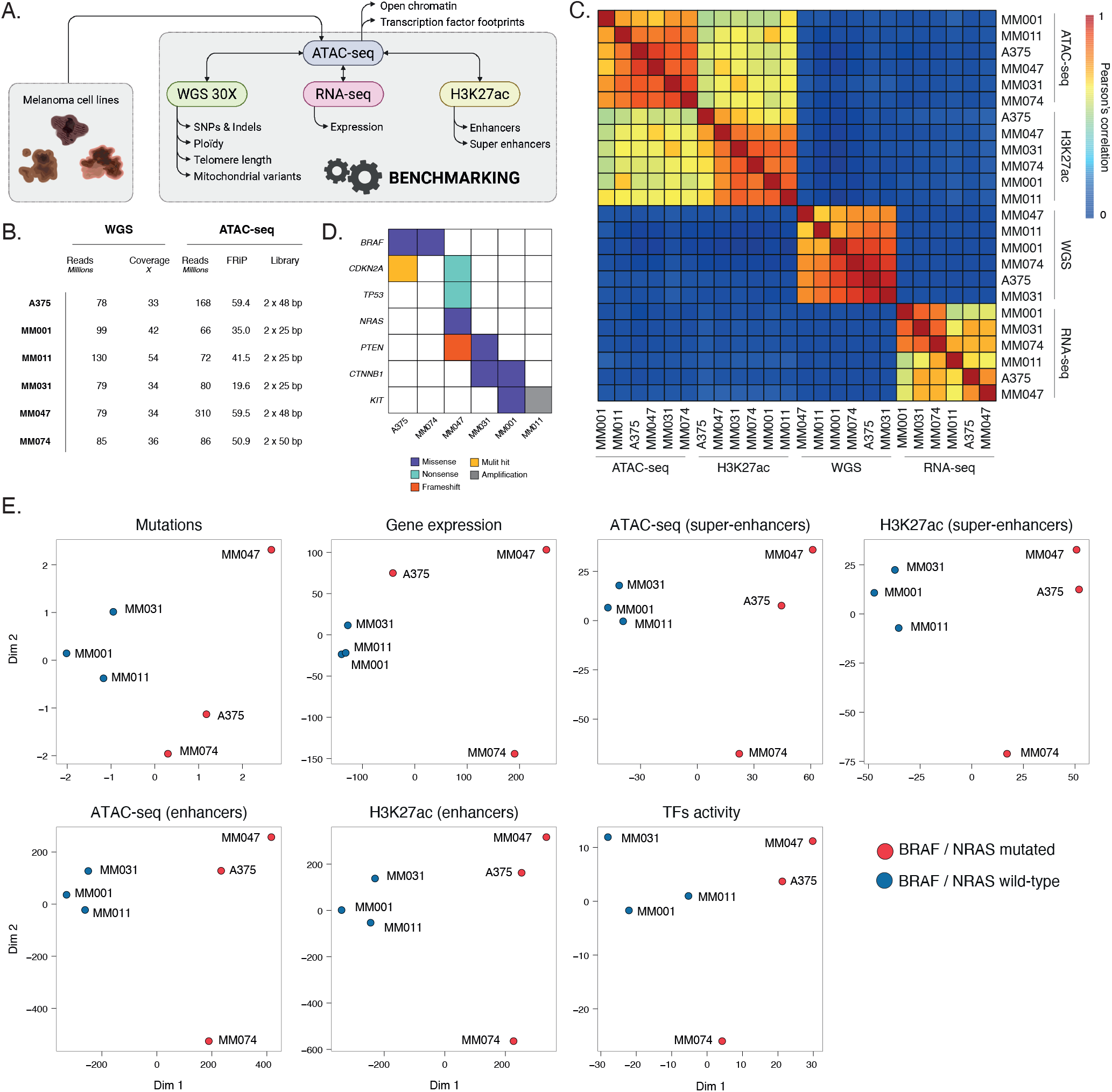
ATAC-seq links oncogenic alterations to coherent regulatory programs in patient-derived melanoma cell lines. **(A)** Study design. Matched bulk WGS, bulk RNA-seq, bulk H3K27ac ChIP-seq, and bulk ATAC-seq were analyzed across 10 patient-derived melanoma cell lines to benchmark ATAC-seq for extracting both genomic and epigenomic features. **(B)** Sequencing metrics for each patient-derived melanoma cell line, including WGS coverage, ATAC-seq fragment yield, and ATAC-seq fraction of reads in peaks (FRiP). For benchmarking, only cell lines with WGS coverage ≥ 30× and ATAC-seq depth ≥ 50 million fragments (A375, MM001, MM011, MM031, MM047 and MM074) were retained. To mitigate confounding effects of heterogeneous ATAC-seq read lengths across datasets, ATAC-seq fragments were hard-trimmed to 2 × 25 bp. **(C)** Genome-wide pairwise Pearson correlation of signal intensity across assays (RNA-seq, ATAC-seq, H3K27ac ChIP-seq and 30× WGS), computed on 1 kb genomic bins and displayed as a clustered heatmap. Rows and columns were hierarchically clustered using correlation distance and Ward’s D^2^ linkage. **(D)** Oncoplot summarizing recurrent melanoma driver alterations (e.g., *BRAF, TP53*, and *NRAS*). across the six benchmarked patient-derived melanoma cell lines. Alterations are annotated by functional class (missense, amplification, frameshift, and multi-hit). **(E)** Block partial least squares (block-PLS) integrative analysis combining mutation data, transcription factor footprinting, chromatin accessibility (in enhancers and super-enhancers), histone modifications (H3K27ac signal at enhancers and super-enhancers), and RNA expression profiles. The block PLS integration was performed in regression mode, with genetic driver alterations data treated as an illustrative outcome.

At the epigenomic level, ATAC-seq accessibility profiles closely mirrored H3K27ac enrichment at H3K27ac-defined regulatory elements and the resulting similarity structure grouped profiles by cell line instead of separating them by modality, showing strong concordance between the two assays at active regulatory regions (Fig. S1). At a broader scale, genome-wide pairwise Pearson correlations computed on 1 kb genomic bins organized datasets mainly by assay and yielded a clustered structure in which ATAC-seq and H3K27ac formed neighboring blocks with higher inter-assay similarity than either showed with RNA-seq or 30× WGS (Fig. 1C). Together, these analyses show that bulk ATAC-seq retains a stable epigenomic signal - at regulatory elements and genome-wide - even under stringent read-length harmonization.

WGS-derived oncogenic driver profiles stratified the patient-derived melanoma cell lines into BRAF/NRAS-mutated versus BRAF/NRAS-wild-type groups (Fig. 1D). To assess whether these genomic alterations are reflected in coordinated regulatory programs, we performed an unsupervised block partial least-squares (PLS) integration in regression mode, integrating gene expression (RNA-seq), chromatin accessibility at enhancers and super-enhancers, H3K27ac signals at enhancers and super-enhancers, and transcription factor activity inferred from ATAC-seq digital footprinting, and treating the WGS driver block as an illustrative outcome (Fig. 1E). This integrative analysis recapitulated the separation between BRAF/NRAS-mutated versus wild-type patient-derived melanoma cell lines, linking oncogenic alterations to coherent changes across chromatin accessibility, histone acetylation, transcription factor activity, and transcriptional output.

These results show that chromatin accessibility captured by bulk ATAC-seq is not only concordant with transcriptional and H3K27ac landscapes, but is structured in a manner that reflects underlying oncogenic alterations, linking regulatory state to genetic context in patient-derived melanoma cell lines. This provides a solid foundation for the subsequent analyses, in which we examine how far genome-derived information can be extracted from the same ATAC-seq libraries.

### ATAC-seq delivers high-confidence SNPs within accessible chromatin in patient-derived melanoma cell lines

We next evaluated bulk ATAC-seq as a substrate for small-variant detection using matched 30× WGS as the reference. As expected for an accessibility-biased assay, comparisons showed that variant overlap between ATAC-seq and WGS outside ATAC-seq peaks was negligible (Table 2, average recall < 6% for SNPs detected with bcftool/mpileup), reflecting the scarcity of allelic sampling in closed chromatin. These results motivated our analysis, which focuses explicitly on small variants located within ATAC-seq peaks, i.e. genomic regions where ATAC-seq provides sufficient and reproducible coverage for genotyping. Within this accessible space, we benchmarked ATAC-seq callsets obtained at increasing sequencing depth (10M–50M fragments) against matched 30× WGS across the six melanoma cell lines (Fig. 2A–B; Table 2).

**Figure 2.**
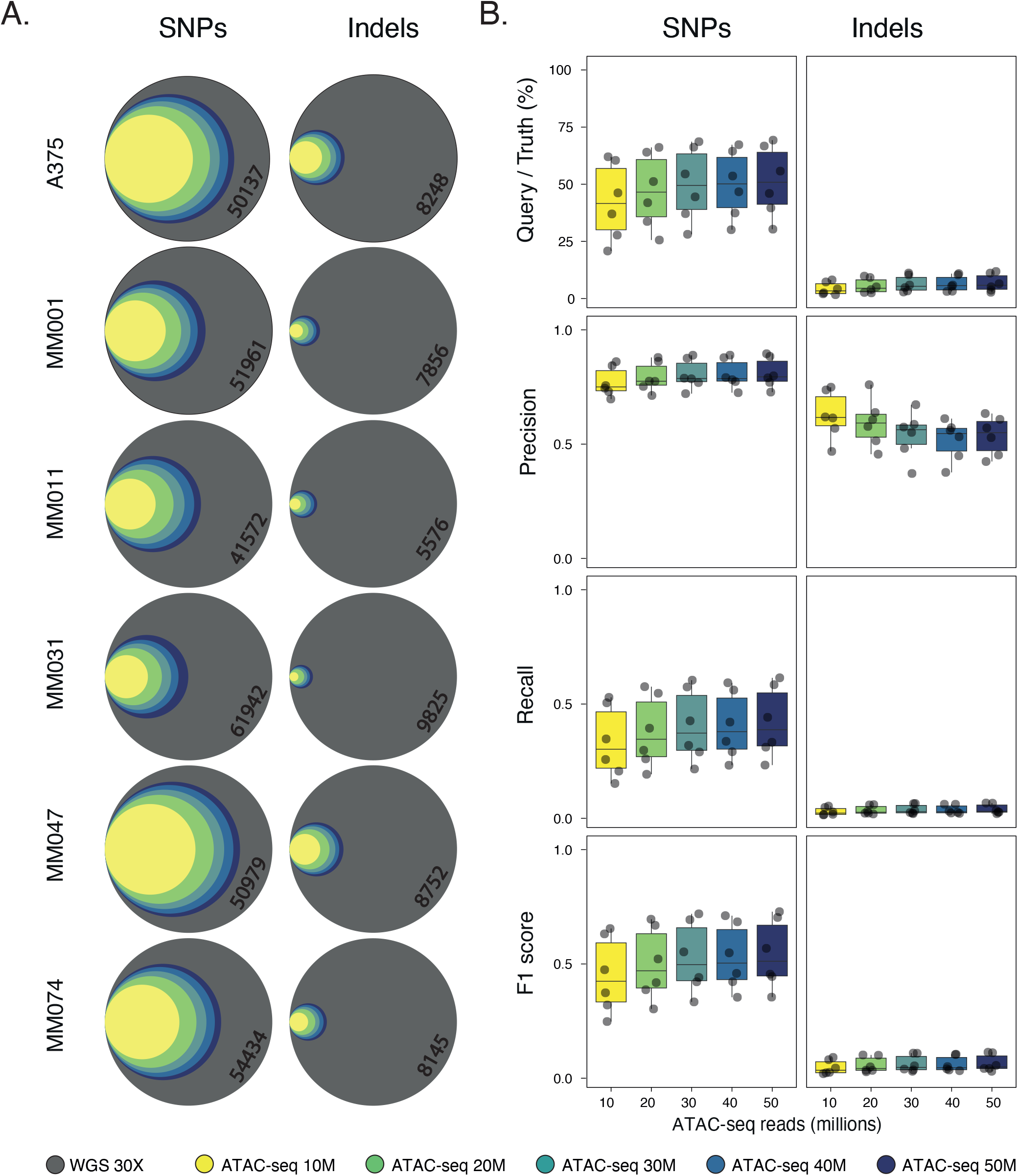
ATAC-seq delivers high-confidence SNPs within accessible chromatin in patient-derived melanoma cell lines. **(A)** Proportional Venn diagrams illustrating the overlap of SNPs and short indels detected with bcftools/mpileup from ATAC-seq within peaks compared with matched 30× WGS for each patient-derived melanoma cell lines. Concentric circles represent ATAC-seq callsets obtained at increasing sequencing depths (10M-50M fragments), while the outer grey circle represents the 30× WGS reference callset. Variants detected exclusively by only one technology across all depths were omitted to improve readability. The number displayed inside the grey circle corresponds to the total number of 30× WGS variants falling within the ATAC-seq accessible space, defined here as the peak set obtained with 50M ATAC-seq fragments. **(B)** Boxplots illustrating the distribution of performance metrics for bcftools/mpileup small-variant calls from ATAC-seq across patient-derived melanoma cell lines at each ATAC-seq sequencing depth (10M-50M fragments). Metrics include percentage overlap with the matched 30× WGS callset, precision, recall and F1 score, computed using ATAC-seq calls as the query and 30× WGS as the truth set, and restricted to variants within ATAC-seq peaks.

As illustrated by positional overlap (Fig. 2A), WGS consistently identified a larger number of variants overall, whereas ATAC-seq recovered a depth-dependent subset enriched in accessible chromatin. SNP recovery improved monotonically with sequencing depth, with the median fraction of WGS SNPs recovered within peaks increasing from ∼40% at 10M fragments to ∼45% at 30M and reaching ∼50% at 50M fragments (Fig. 2B, Table 2). Beyond ∼30–40M fragments, gains in overlap were modest, indicating a clear diminishing-returns regime. Importantly, at the genotype level, ATAC-seq-derived SNPs were characterized by consistently high precision across all depths, with median precision values exceeding 0.75 at 10M fragments and slightly increasing up to ∼0.80–0.85 at higher sequencing depth (Fig. 2B). In contrast, recall remained the limiting factor, with median recall values below 0.40 at all depths (typically ∼0.25–0.30 at 10M and ∼0.35–0.40 at 50M). As a result, F1 scores increased with depth (from ∼0.40 at 10M to ∼0.55 at 50M) but remained bounded by recall rather than precision (Fig. 2B). This pattern reflects a regime in which ATAC-seq preferentially recovers a high-confidence subset of variants, while incomplete allelic sampling remains the dominant source of discordance with WGS.

Indel detection further highlighted the impact of local sequence context and read length on variant recovery. In this first benchmark dataset, where ATAC-seq reads were harmonized to 2 × 25 bp to control for library heterogeneity, indel overlap between ATAC-seq and WGS remained very limited across all depths (<10%), with median recall values below 0.05 even at 50M fragments (Fig. 2B, Table 2). Precision was moderate (typically ∼0.4–0.6, depending on depth), but F1 scores remained uniformly low (<0.1), reflecting a severe sensitivity bottleneck. This behavior is consistent with insufficient local sequence context for accurate alignment and breakpoint resolution under short-read conditions and therefore might reflects a read-length limitation rather than a fundamental constraint of accessibility-based sampling. We explicitly examine the impact of longer read length in subsequent analyses of the LGG/GBM cohort.

To assess the robustness of these observations to variant-caller choice, we compared bcftools/mpileup, FreeBayes, DeepVariant, and GATK HaplotypeCaller (Fig. S2A-B, Table 2). For SNPs, bcftools/mpileup consistently achieved the highest recovery, with median overlap increasing from ∼40% at 10M to ∼50% at 50M fragments and recall rising from ∼0.25 to ∼0.35. FreeBayes showed intermediate performance (median overlap increasing from 30% to 42%; median recall from 0.22 to 0.34), whereas DeepVariant and HaplotypeCaller recovered fewer SNPs overall (median overlap typically ≤35% at 50M, and median recall typically <0.32). Precision differences across callers were comparatively modest, generally ranging from ∼0.80 to ∼0.93, resulting in the highest SNP F1 scores for bcftools/mpileup at higher depth (Fig. S2B). In contrast, indel performance remained uniformly limited across callers, with recall typically in the 0.01–0.04 range at 50M fragments and F1 scores below 0.1 irrespective of the method used. Overall, bcftools/mpileup provided the best trade-off for ATAC-seq-derived calls in this context, maximizing SNP recovery (highest overlap, highest recall and F1 at high depth) while maintaining significant precision. Indel detection remained comparatively limited across callers (Fig. S2B).

Mechanistically, variant recoverability closely tracked local ATAC-seq sampling. SNPs shared between ATAC-seq and 30× WGS were supported by substantially higher ATAC-seq coverage, with median fragment counts increasing from ∼7 fragments at 10M to ∼18 fragments at 50M, whereas WGS-only SNPs typically remained supported by fewer than five ATAC-seq fragments even at higher depth (Fig. S2C). Shared SNPs were also preferentially located near ATAC-seq peak summits (median distance ∼100 bp), whereas WGS-only SNPs were located in genomic regions further away from the corresponding peak centers (median distance ∼500 bp), consistent with reduced allelic sampling away from regions of maximal accessibility. Indels followed similar trends but with stronger attrition in recall, reinforcing that in bulk ATAC-seq, depth helps, yet effective local sampling within peaks - especially near summits - remains a key determinant of variant recovery.

Together, these results demonstrate that bulk ATAC-seq supports high-confidence SNP genotyping within accessible chromatin, operating in a regime of high precision and depth-dependent but intrinsically limited recall. Rather than approximating genome-wide variant discovery, ATAC-seq provides a high-confidence genotyping readout precisely anchored in regulatory DNA, enabling direct interrogation of sequence variation within functionally active chromatin.

### ATAC-seq recapitulates WGS copy-number alterations in patient-derived melanoma cell lines

We then asked whether bulk ATAC-seq could recover large-scale copy-number variation by leveraging background coverage - *i*.*e*. genome-wide signal outside relaxed ATAC-seq peaks - to minimize fluctuations driven by accessibility in regulatory elements. Using QDNAseq at a 100 kb resolution, ATAC-seq-derived log_2_ copy-number profiles closely tracked matched 30× WGS across all six patient-derived melanoma lines and across downsampling levels (10M-50M fragments, Fig. 3A). In per-bin comparisons, distinct copy-number regimes were already apparent at modest depth, with clearly delineated clouds corresponding to diploid bins (2N), losses/LoH states (≤1N), and amplifications (≥3N), and agreement tightening progressively as depth increased (Fig. 3A). Consistent with this depth-dependent sharpening, genome-wide concordance improved modestly and stabilized early (Fig. 3B), with median correlations of ∼0.66 at 10M fragments, ∼0.69 at 20M, and ∼0.70 from 30M to 50M., indicating that most CNV structure can be recovered without requiring the deepest ATAC-seq libraries (Fig. 3B).

**Figure 3.**
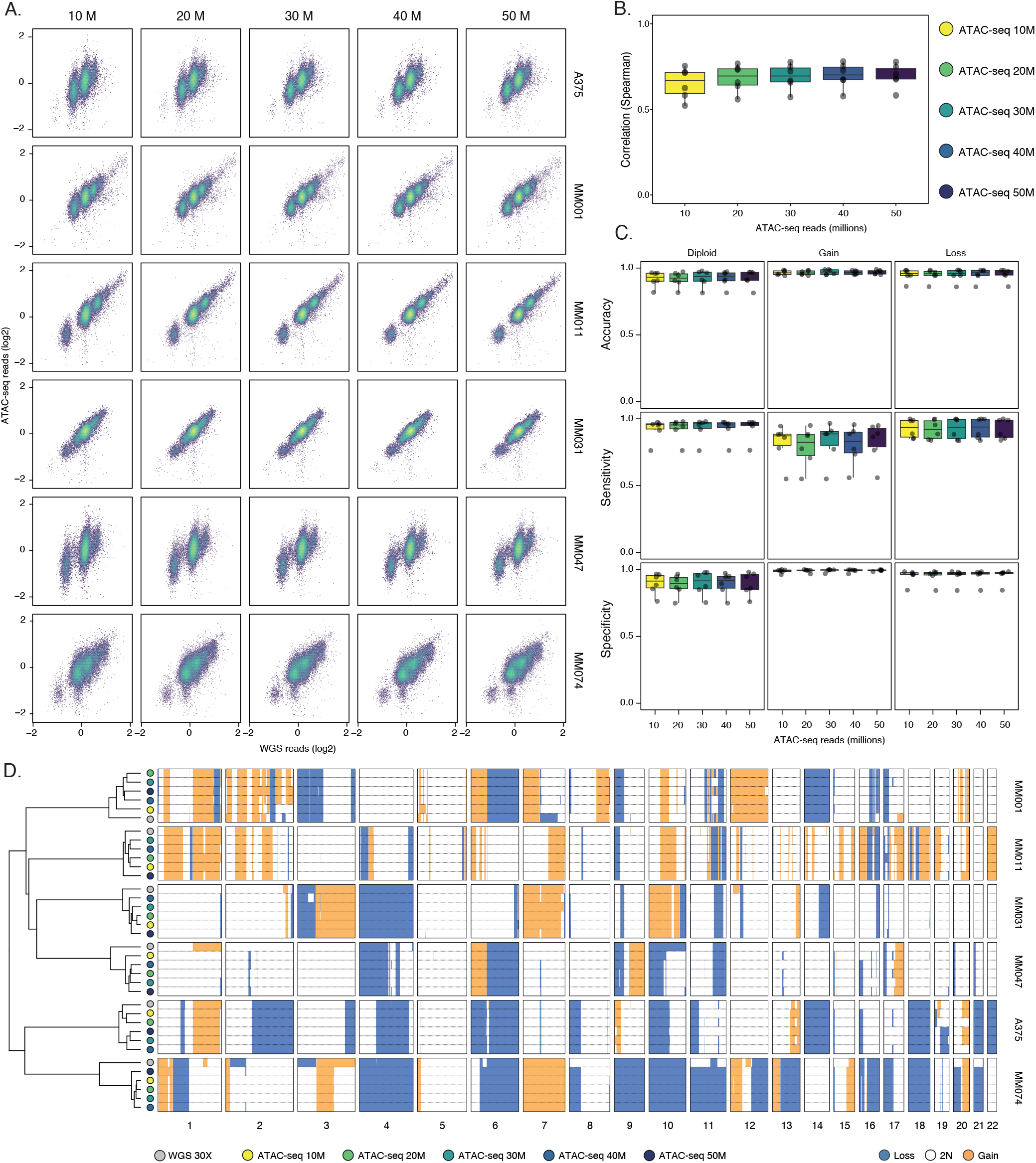
ATAC-seq recapitulates WGS copy-number alterations in patient-derived melanoma cell lines. **(A)** Hexbin scatterplots comparing log_2_-transformed copy-number estimates in 100 kb genomic bins derived from 30× WGS and from ATAC-seq at different sequencing depths (10M-50M fragments) for each patient-derived melanoma cell lines. **(B)** Boxplot illustrating the distribution of Spearman correlation coefficients between ATAC-seq and 30× WGS copy-number estimates in 100 kb genomic bins across patient-derived melanoma cell lines as a function of ATAC-seq depth. **(C)** Boxplots illustrating the classification performance of ATAC-seq copy-number calls relative to matched 30× WGS across patient-derived melanoma cell lines summarized by accuracy, sensitivity, and specificity and stratified by class (loss, diploid, gain). **(D)** Genome-wide CNV profiles inferred from 30× WGS and ATAC-seq at different sequencing depths (10M-50M fragments) for each patient-derived melanoma cell line. Samples were clustered using Pearson correlation and Ward’s D^2^ linkage. Copy-number gains are highlighted in orange, losses in blue, and diploid segments in white.

To move beyond correlation, we classified bins into loss (≤1N), diploid (2N), or gain (≥3N) and compared ATAC-seq states to 30× WGS as a reference (Fig. 3C). Across depths, accuracy remained high for all classes (>0.9 for diploid; >0.95 for gains and losses), and CNV calling behaved conservatively: specificity for both gains and losses stayed near ceiling (>0.95), indicating that ATAC-seq rarely introduces spurious non-diploid states relative to 30× WGS. Depth barely improved sensitivity, and revealed a clear class dependence - diploid bins being most consistently recovered, losses generally robust, and gains showing lower sensitivity and wider dispersion, consistent with greater dependence on segmentation thresholds and coverage uniformity (Fig. 3C). Consistently, genome-wide CNV profiles were highly stable within each patient-derived melanoma cell line across downsampling. Hierarchical clustering of genome-wide CNV profiles grouped samples primarily by cell line, with ATAC-seq profiles remaining close to their matched 30× WGS counterparts even at reduced depth (Fig. 3D), and genome-wide segmentation indicated that major arm-level patterns as well as representative focal events were retained across depths (such as the *cKIT* amplification in MM011, Fig. S3).

These results establish bulk ATAC-seq as a reliable and depth-efficient readout of large-scale copy-number architecture, capturing genome structure with high specificity.

### ATAC-seq reads out canonical telomeric repeats abundance in patient-derived melanoma cell lines

We next examined whether bulk ATAC-seq retains measurable signal over telomere-associated repeats in the patient-derived melanoma cell lines. Telomeric repeat abundance was quantified using TelomereHunter across all autosomal chromosome ends and compared between ATAC-seq libraries downsampled from 10M to 50M fragments and matched 30× WGS (Fig. 4A–B). To reduce confounding contributions from intrachromosomal and interstitial telomeric repeats, downstream analyses were restricted to intratelomeric read counts.

**Figure 4.**
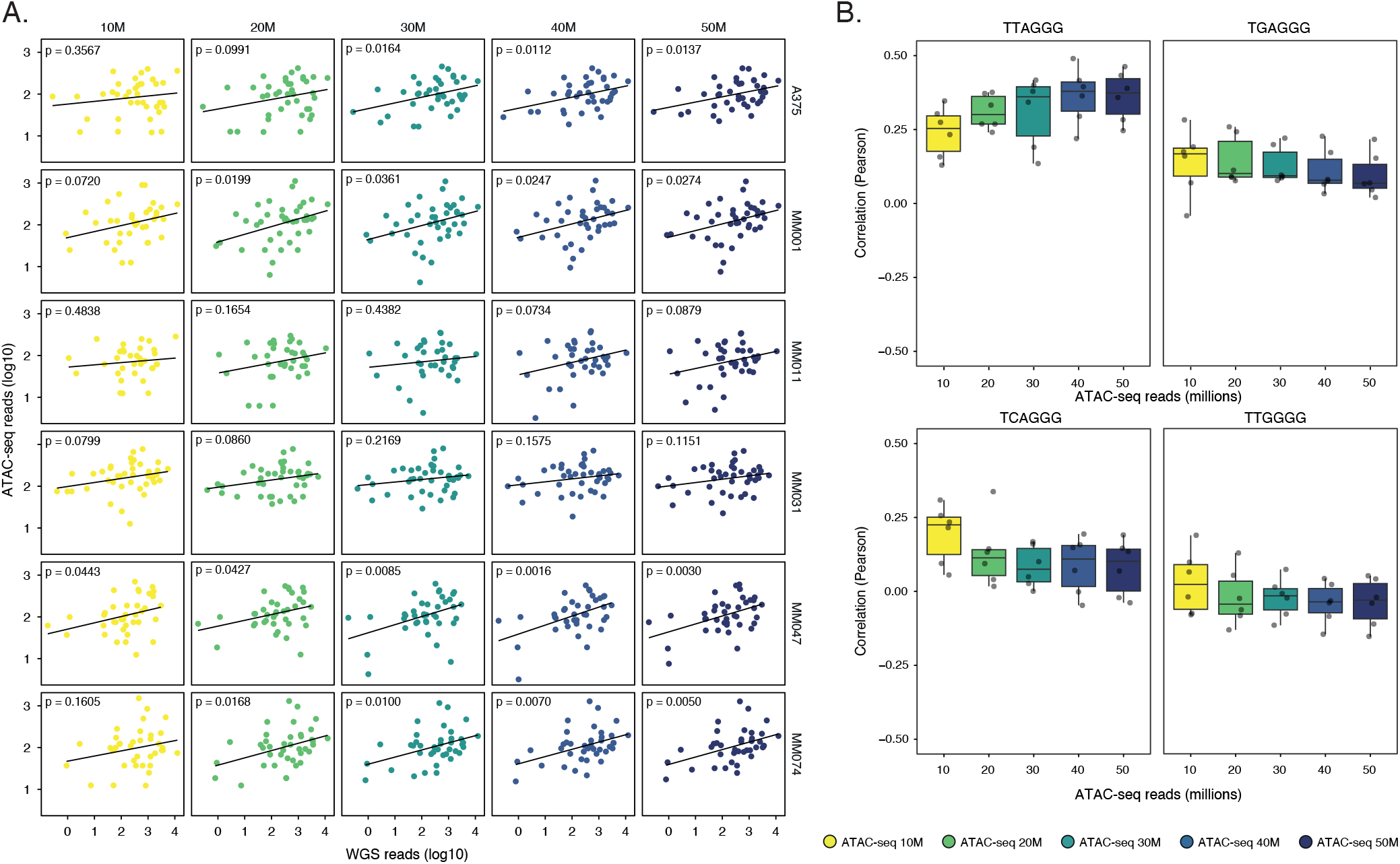
ATAC-seq reads out canonical telomeric repeats abundance in patient-derived melanoma cell lines. **(A)** Scatterplots of log_10_-transformed telomeric repeat content estimated from 30× WGS and ATAC-seq at different sequencing depths (10M-50M fragments) for each patient-derived melanoma cell line. Each dot represents the log_10_-transformed telomeric repeat content quantified at one of the 2 x 22 autosomal telomere for the canonical TTAGGG telomeric motif. The black line indicates the least-squares regression fit. Pearson correlation test *p*-values are indicated in each panel. **(B)** Boxplot illustrating the distribution of Pearson correlation coefficients between ATAC-seq and 30× WGS telomeric repeat content across patient-derived melanoma cell lines, shown as a function of ATAC-seq sequencing depth. The analysis is stratified by telomeric repeat sequence (TTAGGG: canonical telomeric repeat sequence).

TTAGGG was the only telomeric motif to display a robust and interpretable depth-dependent behavior in this cohort (Fig. 4A–B). At the individual cell-line level, concordance with 30× WGS varied across samples but consistently improved with sequencing depth. In A375, correlations were non-significant at 10–20M fragments and became significant at ≥30M (p = 0.0164), whereas MM047 already showed significant concordance at 10M (p = 0.0443), which further strengthened with depth (p = 0.0016 at 40M, Fig. 4A). When aggregated across cell lines, TTAGGG correlations increased from a median of 0.25 at 10M to 0.38 at ≥30M and then plateaued (Fig. 4B), indicating that additional sequencing primarily refines the precision of repeat-abundance estimates rather than revealing qualitatively new signal. In contrast, telomeric variant repeats (TGAGGG, TCAGGG and TTGGGG) showed a negative concordance and a weaker, non-monotonic response to sequencing depth (Fig. 4B). The specificity of the depth effect to TTAGGG argues against a uniform technical gain and instead mirrors telomere sequence organization: long distal telomeric tracts are dominated by TTAGGG, whereas variant repeats are preferentially enriched near the proximal telomere-subtelomere junction and are therefore less uniformly represented across chromosome ends (65). Under this architecture, increasing ATAC-seq depth is expected to stabilize quantification of the most abundant motif before it meaningfully improves estimates of rarer, context-dependent variants - an imbalance that is likely accentuated in this cohort by the standardized 2 × 25 bp read-length harmonization.

Importantly, recent work has clarified that telomere-like reads in ATAC-seq cannot be interpreted as a direct proxy for telomere length, as a substantial fraction originates from subtelomeric regions and may reflect chromatin condensation or accessibility states rather than physical telomeric tracts (35, 65). In line with this, we deliberately constrained our analysis to telomere-associated repeat abundance rather than length per se. To further reduce confounding from intrachromosomal pseudo- and interstitial telomeric repeats, TelomereHunter outputs were restricted to intratelomeric read counts, thereby focusing on the subset of repeat signal most consistently associated with chromosome ends. Within this framework, ATAC-seq captures a reproducible telomere-associated repeat signal - most robustly for the canonical TTAGGG motif - whose behaviour is internally coherent across depth and samples. We extended this analysis to the TCGA LGG/GBM cohort, where larger cohort size enabled testing the association between ATAC-seq-derived telomere-associated repeats and *TERT* expression.

### ATAC-seq enables high-coverage mitochondrial profiling in patient-derived melanoma cell lines

Next, we asked whether bulk ATAC-seq can be leveraged for mitochondrial genome profiling and whether the depth-efficient sampling of mtDNA translates into robust, quantitative mitochondrial allele-fraction fingerprints. Because mitochondrial genomes are present in a high copy number and are not packaged into canonical nucleosomes, they constitute an exceptionally efficient substrate for Tn5 transposition. Consistently, mitochondrial coverage in ATAC-seq was exceptionally high and increased with sequencing depth (Fig. 5A) and reached ∼10,000× mitogenome coverage already at 10M fragments for most of the patient-derived melanoma cell lines, revealing mitochondrial DNA as one of the most efficiently sampled genomic compartments in bulk ATAC-seq libraries.

**Figure 5.**
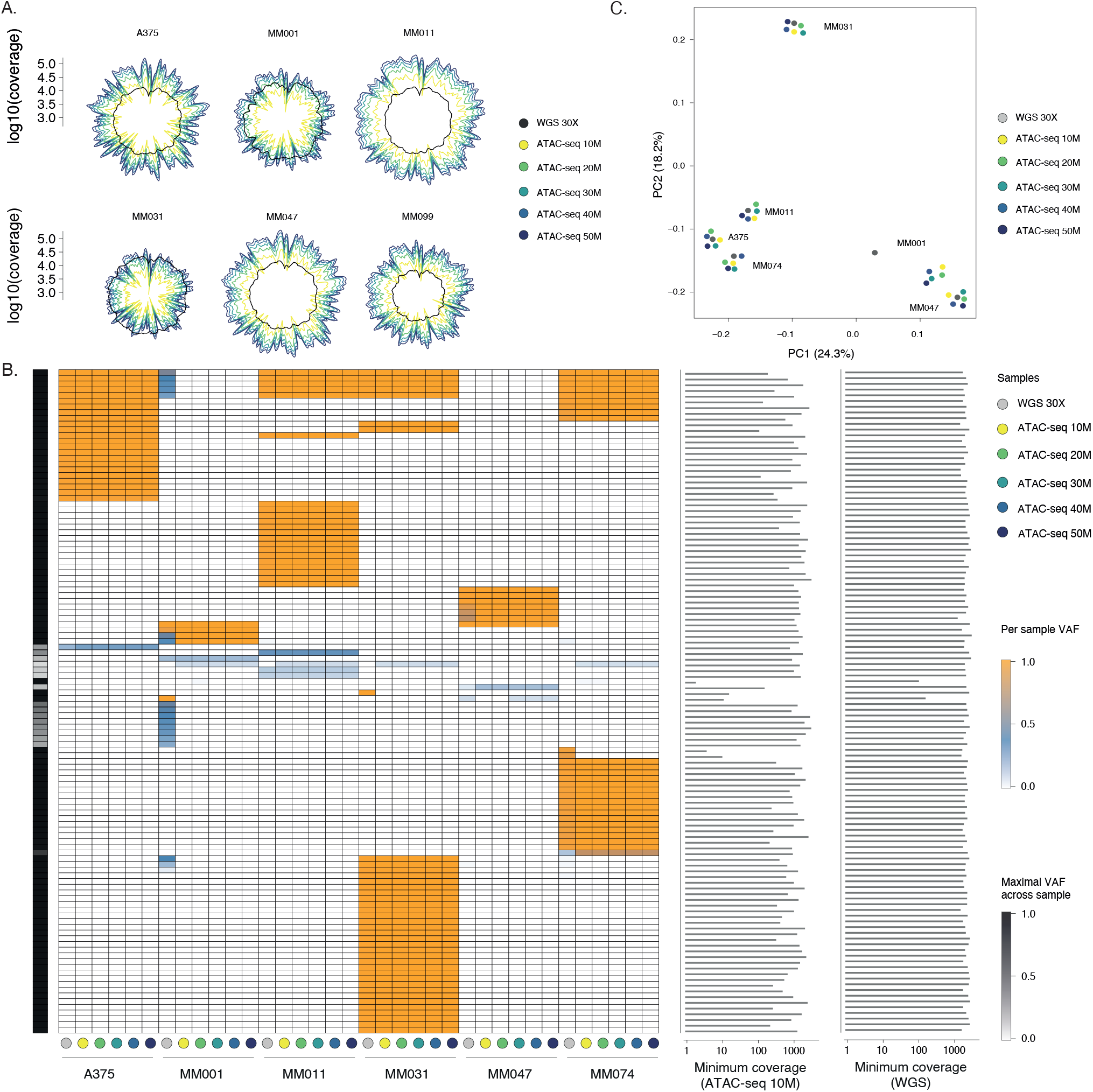
ATAC-seq enables high-coverage mitochondrial profiling in patient-derived melanoma cell lines. **(A)** Polar representation of log_10_-transformed mitogenome coverage for each patient-derived melanoma cell line comparing 30× WGS with ATAC-seq at increasing sequencing depths (10M-50M fragments. **(B)** Heatmap of mitochondrial SNP variant allele frequencies (VAFs) for variable sites with standard deviation of VAF > 0.1 across samples. Rows correspond to SNPs and columns to samples across platforms (30× WGS and ATAC-seq) and ATAC-seq depths. Hierarchical clustering uses Pearson correlation and Ward’s D^2^ linkage. Fixed variants (VAF = 1) are shown in orange, whereas subclonal or heteroplasmic variants (VAF < 1) are shown in blue shades. The left-side annotation bar reports the maximal VAF observed for each SNP across all samples. Two annotation histograms on the right display, for each SNP, the minimum coverage observed across patient-derived melanoma cell lines with 10M ATAC-seq fragments and with 30× WGS, respectively, allowing assessment of coverage-dependent SNP detection. **(C)** Principal component analysis (PCA) of mitochondrial SNP VAF profiles at variable sites with standard deviation of VAF > 0.1. For each patient-derived melanoma cell line, one dot corresponds to the 30× WGS dataset (grey), while additional points correspond to ATAC-seq datasets downsampled to increasing sequencing depths (10M-50M fragments, yellow-to-dark-blue gradient). The first two principal components explain 24.3 + 18.2 = 42.5% of the total variance.

To enable direct cross-platform and cross-depth comparisons, we used mgatk in bulk mode and performed joint calling across all libraries, treating each downsampled ATAC-seq dataset (10M–50M) and the matched 30× WGS as an independent pseudo-sample within a single callset. This design ensures that variant allele fractions (VAFs) are computed over the same catalogue of candidate sites. After stringent filtering aimed at removing artefact-prone positions - based on strand concordance (forward vs reverse counts), variance-to-mean ratio (VMR), and coverage constraints - we retained 284 high-confidence mtDNA SNPs for downstream analyses. The resulting VAF heatmaps revealed highly structured, cell line-specific mtDNA allele-fraction fingerprints that were preserved across ATAC-seq downsampling and closely matched the corresponding 30× WGS profiles (Fig. 5B). Within each cell line, ATAC-seq profiles remained coherent from 10M to 50M fragments and aligned with 30× WGS, producing stable blocks of variants near fixation (VAF ≈ 1) together with a smaller subset of reproducibly sub-fixed sites. Importantly, because these are bulk profiles, VAF<1 should be interpreted as population-level allele mixtures, integrating intracellular heteroplasmy and/or mixtures of subclones, rather than as per-cell heteroplasmy. Across cell lines, residual ATAC–WGS discrepancies were confined to a small subset of sites and were preferentially observed at loci with lower mitochondrial coverage in the shallowest ATAC-seq libraries (i.e. 10M fragments; Fig. 5C). This pattern supports local under-sampling as the dominant source of disagreement, rather than systematic platform-specific shifts in VAF estimates (Fig. 5C)

Global similarity across samples was confirmed by PCA on variable-site VAF profiles: ATAC-seq points clustered close to their matched 30× WGS sample even at 10M fragments, while separating cell lines along the major axes of variation (PC1 = 24.3%, PC2 = 18.2%; 42.5% total variance, Fig. 5C). Together, these analyses show that, in patient-derived melanoma cell lines, bulk ATAC-seq provides not only deep mtDNA coverage but also stable, depth-robust mitochondrial allele-fraction profiles that closely track WGS, establishing mtDNA profiling as one of the most reproducible genome-derived readouts obtainable from standard ATAC-seq libraries.

### ATAC-seq delivers reliable small-variant calls within accessible regions and supports cohort genetic stratification in primary tumors

To evaluate the performance of bulk ATAC-seq for small-variant detection in a clinically relevant context, we extended our benchmark to 22 primary TCGA brain tumors (66, 67) annotated as low-grade gliomas (LGG, n = 13) or glioblastomas (GBM, n = 9), each profiled by bulk ATAC-seq and matched WGS (Fig. 6A; Table 1).This cohort spans a broad clinical range (age 22-70, median 44; 12 males and 10 females) and includes samples collected from multiple anatomical sites (Table 2). Compared with the selected patient-derived melanoma cell line panel, this cohort represents a substantially more challenging setting, combining variable tumor purity, biological heterogeneity, and wide variation in ATAC-seq signal quality (FRiP 8.0-55.1%, mean 27.1 ± 12.2%; Table 1). Of note, ATAC-seq libraries in this cohort were generated with longer read lengths (2 × 76 bp), allowing us to assess how sequencing design modulates variant recovery.

**Figure 6.**
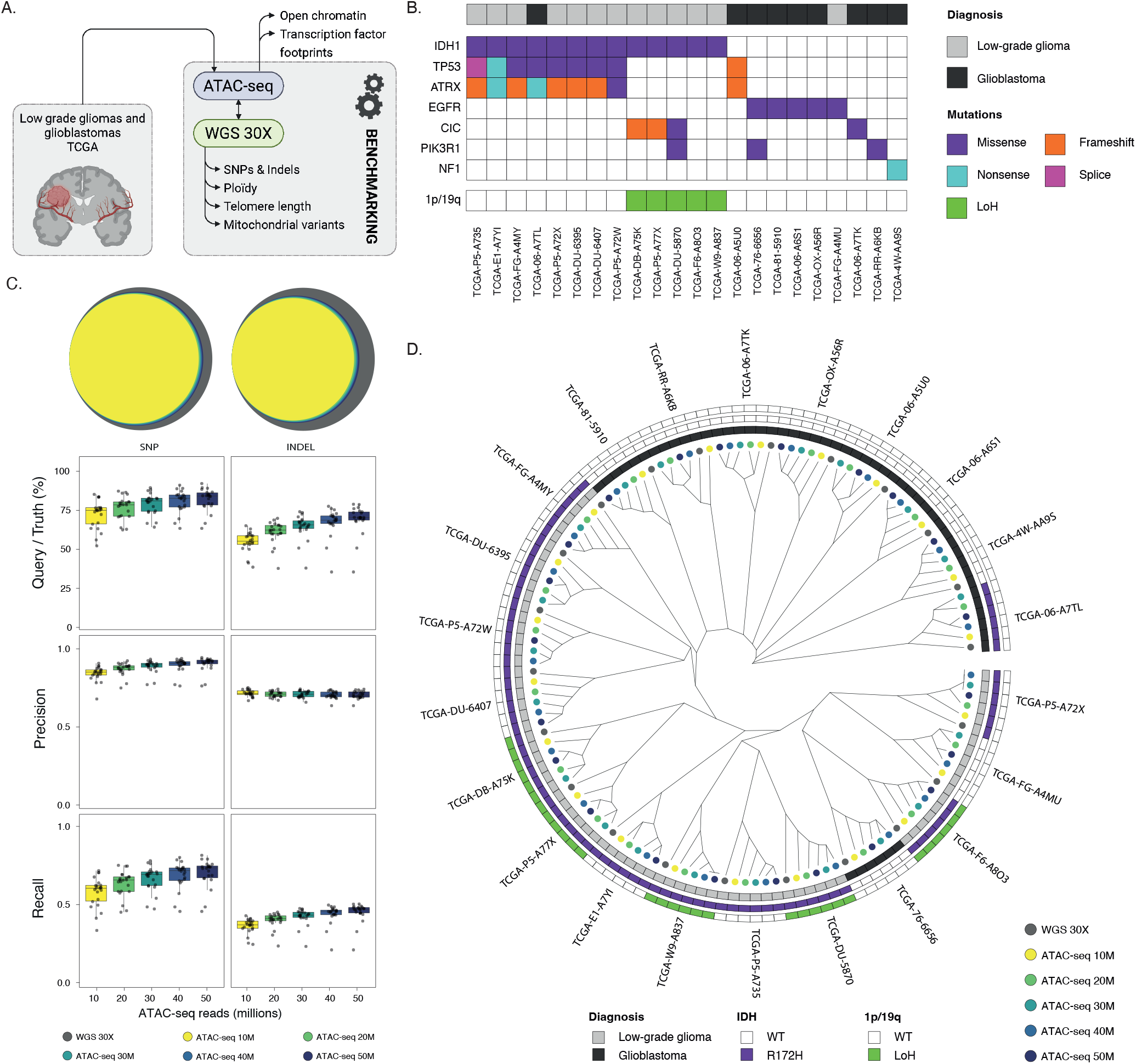
ATAC-seq delivers reliable small-variant calls within accessible regions and supports cohort genetic stratification in primary tumors. **(A)** Study design for the primary tumor cohort comprising 22 TCGA tumors (13 low-grade gliomas and 9 gliomas) with matched bulk ATAC-seq and WGS. **(B)** Oncoplot summarizing recurrent glioblastoma (GBM) and low-grade glioma (LGG) driver alterations across the TCGA cohort (e.g., *IDH1, TP53, ATRX*, 1p/19q). Alterations are annotated by functional class (missense, nonsense, frameshift, splice-site affecting and LoH). **(C)** Boxplots illustrating the distribution of performance metrics for bcftools/mpileup small-variant calls from ATAC-seq TCGA tumors at each ATAC-seq sequencing depth (10M-50M fragments). Metrics include overlap with the matched 30× WGS callset, precision, recall and F1 score, computed using ATAC-seq calls as the query and 30× WGS as the truth set, and restricted to variants within ATAC-seq peaks. **(D)** Phylogenetic reconstruction based on identity-by-state (IBS) genetic distances computed from genotypes at 1,220 SNPs (missing rate < 10% across the global callset), comparing 30× WGS and ATAC-seq callsets generated at increasing sequencing depths (10M-50M fragments) for each TCGA tumor. Outer annotation tracks indicate diagnosis (LGG or GBM), *IDH* status, and 1p/19q status.

Across this cohort, driver alterations followed expected LGG/GBM patterns (Fig. 6B). Most LGG samples carried *IDH1*^R132H^ mutation, frequently co-occurring with *TP53* and *ATRX* alteration (astrocytomas), and a subset displayed 1p/19q loss together with *CIC* mutations (oligodendrogliomas, Fig. 6B). In contrast, GBM samples were enriched for alterations classically associated with aggressive *IDH*-wildtype disease, including *EGFR* and *NF1* point mutations (Fig. 6B). Notably, two samples showed IDH/diagnosis discordance relative to WHO 2021(68) expectations (*IDH1*^R132H^ GBM; *IDH1*-wildtype LGG; Table 2), most plausibly reflecting legacy TCGA annotations and modern molecular reclassification (68).

Despite more variable signal-to-noise ratios compared with the selected patient-derived melanoma cell line panel, bulk ATAC-seq recovered a substantial fraction of 30× WGS-derived small variants within accessible chromatin in the TCGA LGG/GBM cohort, with performance improving steadily as sequencing depth increased (Fig. 6C). For SNPs, median overlap between ATAC-seq and 30× WGS calls increased from ∼75% at 10M fragments to >80% at 50M fragments. Across depths, precision remained consistently high (typically 0.85–0.92), while recall showed a clear depth dependence, rising from ∼0.58 at 10M to ∼0.72 at 50M fragments (Fig. 6C). Indel detection, while consistently more challenging than SNP recovery, showed improved performance in the TCGA LGG/GBM cohort relative to the patient-derived melanoma cell line benchmark. Median indel overlap increased from ∼55% at 10M fragments to ∼70% at 50M fragments, with recall rising from ∼0.36 to ∼0.46 and precision generally remaining above 0.60 across depths (Fig. 6C). This contrasts with the strong recall attrition observed in the patient-derived melanoma cell line analysis after harmonizing reads to 2 × 25 bp and is consistent with the longer 2 × 76 bp ATAC-seq reads in this cohort providing improved local alignment context and breakpoint resolution (Fig. S6).

Beyond feature-level overlap, we next asked whether ATAC-seq-derived genotypes are sufficiently stable to preserve sample identity and cohort structure. Using a set of 1,220 SNPs with low missingness (< 10%) across callsets, we computed identity-by-state (IBS) distances and reconstructed a genotype-based phylogeny across 30× WGS and ATAC-seq datasets at increasing depths (Fig. 6D). Strikingly, for each tumor, ATAC-seq callsets from all downsampling levels clustered tightly with the corresponding 30× WGS genotype, demonstrating high within-sample consistency even at the lowest sequencing depth. Because TCGA does not provide ATAC-seq data from matched adjacent normal tissue for these tumors, the SNP panel used here is expected to capture a mixture of germline SNPs and tumor-derived somatic alterations. Consequently, the higher-level between-sample structure in the IBS tree should primarily be interpreted as reflecting population genetic structure, rather than tumor biology. In this subset, demographic annotations are strongly imbalanced (Table 1), with most GBM tumors derived from patients who self-identified as Black or African Americans, whereas most LGG tumors were derived from patients who self-identified as Whites, so part of the apparent separation between LGG and GBM may reflects underlying germline ancestry and its confounding with cancer type diagnosis.

Together, these results show that bulk ATAC-seq in primary tumors delivers reliable and depth-scalable small-variant calls within accessible chromatin, with high precision and recall in primary tumors. This level of stability supports the use of bulk ATAC-seq to interrogate sequence variation embedded within regulatory elements also enabling cohort-level stratification.

### ATAC-seq captures coordinated CNV, telomere-repeat and mitochondrial signatures in primary tumors

We then examined whether multiple genome-derived signals - spanning genome structure, repetitive elements, and mitochondrial variation - can be recovered in a coordinated manner from the same bulk ATAC-seq libraries generated from primary tumors. Using the TCGA LGG/GBM cohort, we therefore examined CNV landscapes, telomere-associated repeat abundance, and mitogenome profiles in parallel across matched WGS and ATAC-seq datasets.

At the level of large-scale genome structure, ATAC-seq-derived CNV profiles closely recapitulated WGS-derived copy-number states across tumors and sequencing depths (Fig. 7A–B). Focusing on the five LGG tumors annotated by TCGA as 1p/19q codeleted (oligodendrogliomas), ATAC-seq background-coverage profiles robustly reproduced the expected broad-arm losses across all downsampling levels (10M–50M fragments), with highly concordant within-sample profiles and consistent clustering patterns relative to matched 30× WGS (Fig. 7A). Across the full cohort, CNV state classification against the 30× WGS truth set showed high accuracy across diploid, gain, and loss states, generally approaching 0.95–1.0 at all sequencing depths (Fig. 7B). This performance was driven by a markedly conservative behavior for non-diploid calls: specificity for both gains and losses remained near ceiling across depths, indicating that ATAC-seq rarely introduces spurious CNV states relative to WGS. Sensitivity does not improve with depth and showed class-dependent variation, with gains displaying greater dispersion than losses, reflecting the greater sensitivity of copy-number gains to segmentation thresholds and local coverage variability.

**Figure 7.**
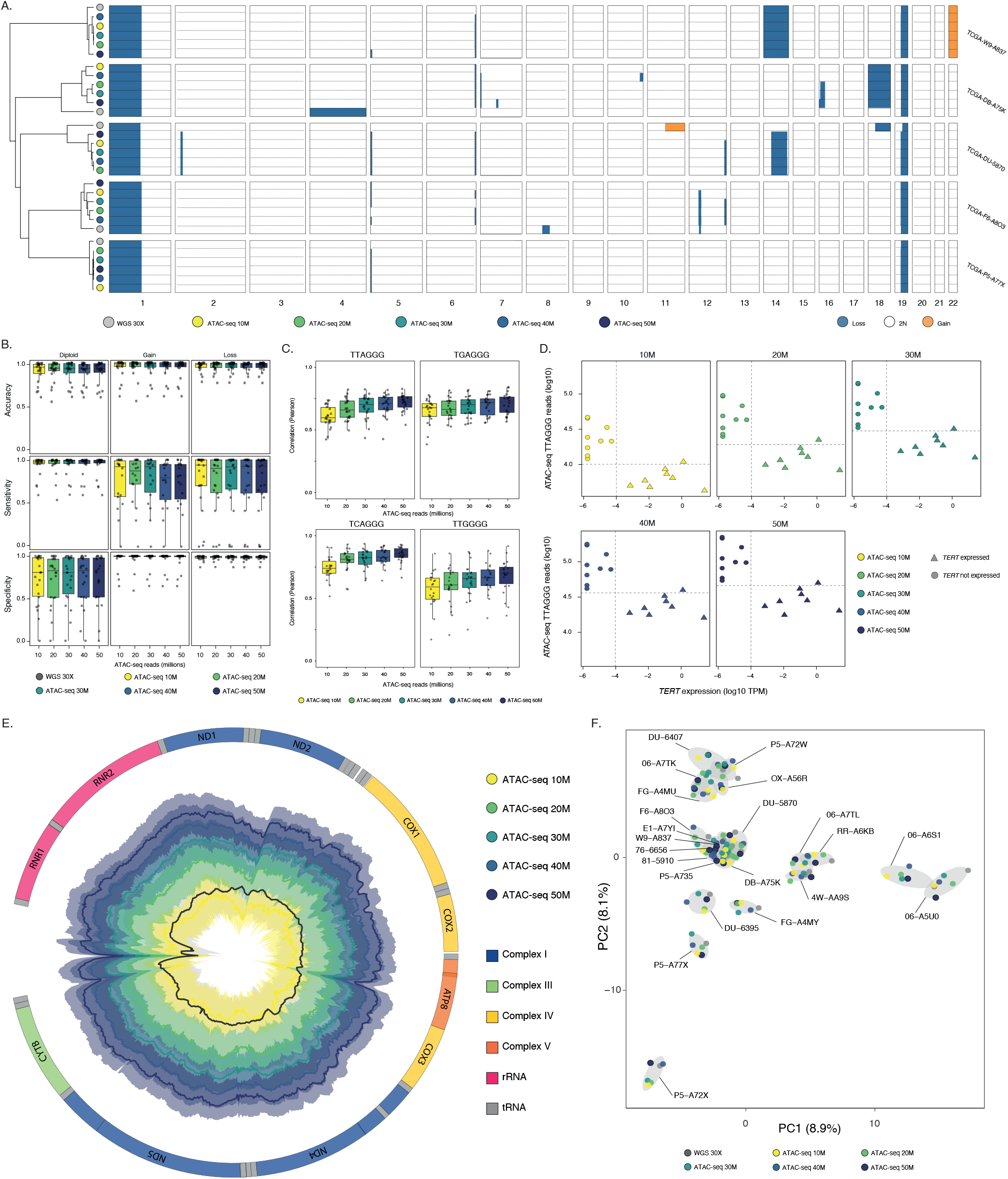
ATAC-seq captures coordinated CNV, telomere-repeat and mitochondrial signatures in primary tumors. **(A)** Genome-wide CNV profiles inferred from 30× WGS and ATAC-seq at different sequencing depths (10M-50M fragments) for the 5 LGG tumors with 1p/19q co-deletion. Samples were clustered using Pearson correlation and Ward’s D^2^ linkage. Copy-number gains are highlighted in orange, losses in blue, and diploid segments in white. **(B)** (A) Boxplots illustrating the classification performance of ATAC-seq copy-number calls relative to matched 30× WGS across LGG and GBM TCGA tumors summarized by accuracy, sensitivity, and specificity and stratified by class (loss, diploid, gain). **(C)** Boxplot illustrating the distribution of Pearson correlation coefficients between ATAC-seq and 30× WGS telomeric repeat content across LGG and GBM TCGA tumors, shown as a function of ATAC-seq sequencing depth. The analysis is stratified by telomeric repeat sequence (TTAGGG: canonical telomeric repeat sequence). **(D)** Relationship between telomeric repeat content and *TERT* expression across TCGA LGG and GBM tumors. Each panel shows ATAC-seq libraries subsampled to increasing sequencing depths (10M-50M fragments). The y-axis indicates log_10_-transformed ATAC-seq reads mapping to the canonical telomeric repeat (TTAGGG), and the x-axis shows log_10_-transformed *TERT* expression (TPM). Points are colored by ATAC-seq depth and shaped according to *TERT* expression status (triangles: *TERT* expressed; circles: *TERT* not expressed). Dashed lines indicate the thresholds used to define high telomeric content and *TERT* expression. **(E)** Polar plot of log_10_-transformed mitogenome coverage across TCGA LGG and GBM tumors. Median profiles are shown for 30× WGS (dark grey) and ATAC-seq at increasing depths (10M-50M fragments, yellow-to-dark-blue gradient). Shaded ribbons indicate the interquartile range (25^th^-75^th^ percentiles) across tumors. The outer ring shows the genomic coordinates of annotated mitochondrial genes along the circularized mitochondrial genome. Gene segments are color-coded according to their assignment to respiratory chain complexes (Complex I, III, IV, V) or RNA genes (rRNA, tRNA). **(F)** Principal component analysis (PCA) of mitochondrial SNP VAF profiles at variable sites with standard deviation of VAF > 0.1. For each LGG and GBM TCGA tumors, one dot corresponds to the 30× WGS dataset (grey), while additional points correspond to ATAC-seq datasets downsampled to increasing sequencing depths (10M-50M fragments, yellow-to-dark-blue gradient). The first two principal components explain 8.9 + 8.1 = 17.0% of the total variance.

In parallel, we evaluated telomere-associated repeat abundance in the same samples and observed that ATAC-seq-WGS correlation distributions shift upward with sequencing depth for all motifs, including the non-canonical repeats (Fig. 7C). In contrast to the patient-derived melanoma cell line cohort where depth-dependent improvements were largely restricted to the canonical TTAGGG motif, primary tumors showed a consistent positive correlation with depth for both canonical and variant telomeric repeats (Fig. 7C). This difference is consistent with the stronger sampling constraint in the melanoma benchmark, where ATAC-seq reads were deliberately hard-trimmed to 2 × 25 bp to harmonize libraries prior to downsampling, thereby reducing local sequence context and effective mappability in repetitive telomeric regions. In the TCGA cohort, the longer 2 × 76 bp reads provide greater anchoring context for repeat-containing fragments, which likely contributes to the more uniform depth-dependent gain in concordance across motifs. Leveraging cohort size within this integrative framework, we then related TTAGGG abundance to *TERT* expression and observed that tumors classified as *TERT*-expressing tended to occupy a lower TTAGGG-repeat regime across depths (Fig. 7D). While counter-intuitive at first glance, this pattern is compatible with a model in which telomerase is reactivated preferentially in contexts of telomere attrition, maintaining telomeres above a minimal functional threshold rather than restoring long telomere tracts, such that high *TERT* expression can coincide with comparatively lower repeat abundance (69).

Finally, ATAC-seq enabled robust mitochondrial profiling in these primary tumor samples. Mitogenome coverage profiles showed that ATAC-seq achieves 30× WGS-like mitochondrial sampling even at modest depth, with increasing depth producing stepwise gains in median coverage across tumors (Fig. 7E). Coverage was broadly distributed along the circular mitogenome but displayed reproducible local dropouts - most notably around *ND2* and *ATP8*, suggesting locus-specific effects that persist across depths (Fig. 7E) which were also captured with 30× WGS. Such structured troughs are consistent with sequence- and context-dependent tagmentation and mapping biases, including Tn5 sequence preference and local mappability constraints. Despite these local features, mitochondrial variant signals were stable: PCA of variable mtDNA SNP VAF profiles maintained tight within-tumor grouping across ATAC-seq depths together with the matched 30× WGS point, indicating that ATAC-seq preserves tumor-specific mtDNA allele-fraction fingerprints in primary samples (Fig. 7F).

Taken together, these analyses show that bulk ATAC-seq in primary tumors captures a coordinated set of genome-derived signatures spanning CNVs, telomere-associated repeats content and mitogenome variations in primary tumors. Importantly, these signals remain internally consistent across sequencing depths and align with matched WGS despite the biological and technical heterogeneity inherent to clinical tumor specimens, reinforcing bulk ATAC-seq as a unified experimental substrate for integrative genome–epigenome characterization in cancer cohorts.

## DISCUSSION

Bulk ATAC-seq was originally conceived as a chromatin accessibility assay, optimized to map regulatory elements and infer transcriptional control across the genome(3, 70). In this study, we show that ATAC-seq also retain reproducible genetic and structural information that extends beyond classical epigenomic readouts. Although much of the methodological development and formal benchmarking of these genome-derived signals has been so far evaluated in single-cell ATAC-seq, where copy-number states, mitochondrial genotypes and genetic subclones can be resolved at cellular resolution (28, 71–73), equivalent reference-grade validation in bulk ATAC-seq has remained comparatively limited. By systematically benchmarking bulk ATAC-seq against matched WGS, we show that ATAC-seq can be used as a multi-layer assay, capturing regulatory state while also supporting selected genomic proxies, provided that each signal is interpreted within its biological and technical limits. To ensure that this benchmark is robust, we structured it around two complementary cohorts. We first analyzed patient-derived melanoma cell lines, where sample composition is controlled and ATAC-seq libraries typically display high signal-to-noise (high FRiP), allowing us to characterize best-case behavior and to isolate technical determinants such as sequencing depth and read length. We then applied the same analyses to 22 primary TCGA brain tumors (LGG/GBM), where ATAC-seq quality metrics vary widely (FRiP range), alongside variable tumor purity and intratumor heterogeneity, thereby more closely reflecting the constraints of clinical material. Together, these two cohorts allow us to define not only what bulk ATAC-seq can recover under controlled conditions, but also which genome-derived signals remain robust and reproducible in heterogeneous tumor samples.

Within this framework, small-variant detection provides a first, stringent test of how far bulk ATAC-seq can be pushed toward genomic readouts. As expected from an accessibility-biased assay, ATAC-seq does not sample the genome uniformly; instead, variant recovery is mostly restricted to accessible chromatin and strongly shaped by local coverage. In the melanoma cell-line cohort, SNP calls derived from ATAC-seq consistently showed high precision across sequencing depths, while recall remained the limiting factor, even within peak regions. This behavior reflects sparse and heterogeneous sampling, and it is reinforced by the strong enrichment of shared SNPs near ATAC-seq peak summits. Importantly, this precision-first regime was generalized to clinically derived samples. In the TCGA LGG/GBM cohort, where ATAC-seq data display longer reads and are biologically heterogeneous, SNP recovery within peaks improved markedly while maintaining high precision across downsampling levels. As a result, ATAC-seq-derived genotypes were stable enough to preserve within-sample identity and to support cohort-scale genetic stratification. These results indicate that, although bulk ATAC-seq cannot provide comprehensive genome-wide variant discovery, it delivers a robust and reproducible genotyping signal within accessible chromatin that is well suited for cohort-level genetic stratification and the analysis of sequence variation occurring within regulatory elements in heterogeneous tumor cohorts. Indel detection further sharpens the distinction between constraints intrinsic to accessibility-based assays and technical limitations imposed by ATAC-seq read length. In the melanoma benchmark, where ATAC-seq reads were harmonized to 2 × 25 bp to control for library heterogeneity, indel recovery was extremely limited across depths, with low recall despite moderate precision. This behaviour is consistent with insufficient local sequence context for accurate alignment and breakpoint resolution (74), and therefore reflects a read-length constraint rather than a fundamental limitation of accessibility-based sampling. In contrast, in the TCGA LGG/GBM cohort profiled with 2 × 76 bp ATAC-seq, indel recovery improved substantially and showed a clear depth dependence, with both overlap and recall increasing across downsampling levels. These results directly link indel performance in bulk ATAC-seq to read length and local alignment context, highlighting that, unlike SNPs, indel detection is particularly sensitive to read length and local alignment context.

Copy-number variation presents a qualitatively different regime. Unlike small variants, CNV inference integrates signal over large genomic windows and is therefore far less sensitive to locus-level sampling sparsity (75). While CNV inference from ATAC-seq has been explored in single-cell settings - where it underpins genetic subclone deconvolution and tumor evolution analyses (30, 72)-its operating characteristics in bulk ATAC-seq have been less systematically evaluated against matched WGS. By exploiting background coverage outside accessible peaks, bulk ATAC-seq robustly recapitulated WGS-derived CNV landscapes across both melanoma cell lines and primary TCGA tumors. Even at modest sequencing depth, ATAC-seq recovered broad-arm alterations and large-scale CNV structure, with concordance stabilizing early as depth increased. Importantly, CNV calling from ATAC-seq behaved conservatively: non-diploid states were detected with high specificity relative to WGS, while sensitivity improved with depth and showed class-dependent variation, with gains showing lower sensitivity and greater variability than losses, consistent with stronger dependence on segmentation thresholds and coverage uniformity. In the TCGA cohort, this conservative behaviour was reinforced at cohort scale, including the robust recovery of canonical alterations such as 1p/19q codeletion in LGG samples. Taken together, these results position CNV profiling as one of the most stable and reliable genome-derived readouts obtainable from bulk ATAC-seq.

One of the most striking genome-derived readouts from bulk ATAC-seq is the mitochondrial genome. In contrast to the nuclear genome, mitochondrial DNA is present in hundreds to thousands of copies per cell and is packaged in nucleoids rather than canonical nucleosomes, making it an exceptionally efficient substrate for Tn5 transposition(3, 4, 32). As with CNVs, the most explicit methodological development and benchmarking of mitochondrial genotyping have been driven by single-cell ATAC-seq(31–33), motivating the need to assess how much of this robustness transfers to bulk ATAC-seq under realistic tumor heterogeneity. As a result, bulk ATAC-seq routinely achieves very high mitochondrial coverage even at modest sequencing depth, often comparable to or exceeding that obtained with matched 30× WGS. In the melanoma cell-line cohort, ATAC-seq reached deep and overall uniform mitogenome coverage already at 10M fragments, enabling stable estimation of mitochondrial allele fractions across a large set of sites. This technical property translates into robust and reproducible mitochondrial variant profiles. Using joint calling across downsampled ATAC-seq libraries and matched WGS, we show that mitochondrial allele-fraction (VAF) patterns are highly consistent across sequencing depths and platforms. In melanoma cell lines, ATAC-seq-derived VAF profiles preserve cell line-specific signatures and closely mirror WGS, with most variants near fixation and a smaller subset of reproducibly sub-fixed sites. Importantly, VAFs < 1 in bulk profiles should not be interpreted as single-cell heteroplasmy, but rather as population-level mixtures reflecting intracellular heteroplasmy and/or subclonal composition. Crucially, this robustness extends to clinically derived samples. In the TCGA LGG/GBM cohort, mitochondrial VAF profiles inferred from ATAC-seq remain highly consistent across downsampling levels and closely match those obtained from the corresponding tumor WGS. This concordance holds despite wide variation in ATAC-seq quality metrics, tumor purity, and biological heterogeneity, indicating that mtDNA allele-fraction patterns are preserved at the sample level and can be reliably recovered from bulk

ATAC-seq. These results highlight mitochondrial profiling as a particularly high-yield and underutilized by-product of bulk ATAC-seq. At the same time, they warrant careful interpretation. Short-read mitochondrial analyses are susceptible to confounding by nuclear mitochondrial DNA segments (NUMTs), which can generate spurious variant calls or distort allele fractions if not properly controlled (76, 77). While our conservative filtering mitigates obvious artefacts (e.g., low-support and strand-inconsistent sites), NUMT-aware mapping and variant-calling strategies remain important avenues for further refinement. Taken together, our results position mitochondrial genome profiling as one of the most robust genome-derived readouts of bulk ATAC-seq, offering reliable and biologically informative signal when interpreted within these technical boundaries.

Telomeric repeats are a fundamentally harder target for bulk ATAC-seq compared to small variants or CNVs. Telomeric regions are indeed highly repetitive, often heterochromatic, and subject to substantial mapping ambiguity, making them intrinsically difficult for short-read sequencing (78–80) and, a fortiori, for an accessibility-based assay. Recent work has emphasized that telomere-like reads in ATAC-seq cannot be straightforwardly interpreted as telomere length, as a substantial fraction originates from subtelomeric domains, interstitial telomeric sequences, or telomere-adjacent repeat tracts, and may reflect chromatin accessibility or condensation states rather than physical telomere tracts (35). Consistent with this view, we deliberately frame our analyses in terms of telomere-associated repeat abundance, and we restrict our readouts to intratelomeric signals to minimize contributions from pseudo- and interstitial repeats. Within this conservative framing, bulk ATAC-seq nonetheless yields reproducible, depth-dependent behavior. In the patient-derived melanoma cell line cohort, where ATAC-seq reads were harmonized to 2 × 25 bp, concordance with WGS-derived repeat abundance was modest and strongly motif-dependent: the canonical TTAGGG signal showed the clearest depth effect, whereas variant repeats remained weaker and less monotonic. This pattern is consistent with limited local context and mappability under short-read conditions, and it underscores that telomeric analyses from ATAC-seq are particularly sensitive to ATAC-seq read length. In the TCGA LGG/GBM cohort profiled with 2 × 76 bp ATAC-seq, correlation distributions shifted upward with depth across motifs, consistent with improved repeat mappability and greater context at longer read length. Leveraging cohort size in the TCGA setting further revealed a biologically coherent association with telomerase activation. Across the TCGA LGG/GBM primary tumors, *TERT*-expressing cases tended to occupy a lower TTAGGG repeat-abundance regime across sequencing depths. While counterintuitive, this observation is compatible with models in which telomerase is preferentially reactivated in the setting of telomere attrition and maintains telomeres above a functional threshold rather than restoring long telomeric tracts (69, 81). Together, these results indicate that telomere-associated repeat abundance can be recovered reproducibly from bulk ATAC-seq under strict constraints. While it does not substitute for dedicated telomere length assays, this signal provides a scalable and comparative readout that can be informative at the cohort level when its scope and limitations are explicitly defined.

This systematic benchmark highlights a simple organizing principle: genome-derived signals recovered from bulk ATAC-seq depend strongly on the scale at which information is integrated. Base-pair-level features such as SNPs operate under tight constraints imposed by local accessibility, allelic balance, and read length, resulting in high precision but limited recall, with indels further exposing the importance of ATAC-seq read length. In contrast, CNV inference integrates signal over large genomic windows and is therefore robust to local sparsity, yielding conservative and stable structural profiles even in heterogeneous tumor samples. Mitochondrial variants occupy a distinct regime, benefiting from high copy number and efficient transposition to produce deep and highly reproducible signal, while telomere-associated repeats represent the most constrained case, where repetitiveness and mapping ambiguity dominate but still allow interpretable cohort-scale trends under strict framing. Taken together, these results show that bulk ATAC-seq does not provide a single level of genomic resolution, but rather a spectrum of recoverable signals whose robustness increases as signal is integrated over broader genomic scales. In practice, this reframes bulk ATAC-seq as an integrative backbone: while regulatory profiling remains the primary readout, the same libraries can also support accessible-region genotyping for quality control and stratification, conservative CNV landscapes for genome-structure context, mitochondrial allele-fraction fingerprints, and cautiously interpreted telomere-associated repeat abundance. The value lies not in competing with WGS, but in enabling a disciplined, multi-layer view of tumor biology from a single assay when multi-platform profiling is constrained by material, cost, or logistics.

Despite this versatility, bulk ATAC-seq has intrinsic limitations that define its boundaries. Its bias toward open chromatin restricts genome-wide variant discovery, leaving deeply heterochromatic and low-accessibility regions underpowered. Indel detection remains particularly sensitive to read length and alignment context, and nucleotide-level analyses are further influenced by sequence-dependent Tn5 transposition bias, and - specifically for mitochondrial calling - by mapping ambiguities such as NUMTs. While Tn5 bias can be modelled and corrected for quantitative coverage-based analyses (17, 78–81), it imposes irreducible limits on qualitative nucleotide-level inference at poorly sampled loci. Similarly, telomere-associated signals require explicit modelling of subtelomeric contributions and benefit from calibration against orthogonal assays. These limitations point toward clear avenues for improvement. Algorithms explicitly designed for ATAC-seq - rather than adapted from WGS - could incorporate accessibility-aware priors for variant calling, chromatin-informed CNV models, NUMT-aware mitochondrial pipelines, and refined repeat-abundance frameworks (32, 33, 73).

In summary, our study demonstrates that bulk ATAC-seq is more than just an epigenomic assay. Our systematic benchmark establishes that a single ATAC-seq library can deliver coordinated regulatory and genomic readouts - capturing chromatin accessibility together with accessible-region genotypes, conservative copy-number architecture, telomere-associated repeat abundance, and mitochondrial allele-fraction fingerprints. This integration confers immediate practical value in settings where tissue availability is limited, multi-platform profiling is impractical, or cost and turnaround time constrain clinical or translational workflows (82–85). Contemporary cancer classification increasingly relies on composite molecular portraits that integrate genetic drivers, copy-number alterations, epigenetic state, telomere maintenance features and metabolic context (85–91). Yet, in practice, these dimensions are rarely interrogated jointly - even in well-resourced precision oncology programs - because they require distinct assays, additional material, and complex analytical coordination. As a result, tumors sharing the same molecular label frequently diverge along axes that are biologically and clinically meaningful. Across multiple cancer types, including diffuse gliomas, melanomas, colorectal cancer and hematological malignancies, variation in genome structure (92– 94), telomere-associated features (64, 65, 95–98), mitochondrial genetics (86, 99) and chromatin accessibility(100, 101) has been linked to treatment response, immune interaction, metabolic reprogramming and evolutionary potential. Despite their recognized importance, these axes are typically studied in isolation, obscuring how they co-evolve within individual tumors and shape integrated tumor behavior (88–96). By extracting these layers from a single assay, genome-aware ATAC-seq enables the definition of composite tumor states that cut across conventional molecular taxonomies. Rather than replacing established classifications, this framework contextualizes them by revealing how regulatory architecture, structural variation, telomere-associated features and mitochondrial genetics co-vary within tumors. Such states provide a structured and biologically interpretable representation of tumor heterogeneity, grounded in measurable molecular features and amenable to retrospective or prospective analysis without additional tissue consumption. Importantly, the value of this approach does not lie in immediate clinical prediction, but in its capacity to organize molecular complexity in a scalable and testable manner (82–85). By maximizing information yield per sample, genome-aware ATAC-seq supports hypothesis-driven exploration of coupled molecular axes that are otherwise fragmented across platforms - offering a pragmatic entry point for integrated tumor profiling when comprehensive multi-omics is infeasible. As precision oncology continues to move toward finer-grained molecular stratification under increasing practical constraints, this integrative use of bulk ATAC-seq provides a versatile and immediately deployable backbone for joint genomic–epigenomic characterization in both research and translational settings.

## SUPPLEMENTARY DATA

The code enabling to reproduce all the analyses performed in this study is hosted on Zenodo with accession 10.5281/zenodo.14871223.

## ACKNOWLEDGEMENTS

We thank Stein Aerts (VIB-KU Leuven Center for Brain and Disease Research, Leuven, Belgium) for kindly providing access to the data and Aurélien de Reynies (Centre de Recherche des Cordeliers, Paris, France) f or helpful discussions and critical feedback. Schematic figures were created in BioRender (Elz, A., 2025).

Author contributions: Islem Toumi (Data curation [equal], Formal analysis [equal], Investigation [equal], Methodology [equal], Software [equal]), Chaimae Kham (Data curation [equal], Formal analysis [equal], Investigation [equal], Methodology [equal], Software [equal]), Lucille Stuani (Data curation [equal], Writing - review & editing [equal], Supervision [equal], Laurent Lecam (Writing - review & editing [equal], Funding acquisition [lead]), and Pierre-François Roux (Conceptualization [lead], Data curation [lead], Formal analysis [lead], Investigation [lead], Methodology [lead], Software [lead], Writing - original draft [lead], Writing - review & editing [lead], Supervision [equal]).

## CONFLICT OF INTEREST

The authors declare no competing interests.

## FUNDING

This work was supported by grants from the Ligue contre le Cancer, the i-SITE Excellence program of Montpellier University under the investissements France 2030, the SIRIC Montpellier Cancer (Grant INCa INSERM DGOS 12553), the ARC foundation, the Institut National contre le Cancer (INCa), the Fondation pour la Recherche Médicale (FRM), and with the institutional support of INSERM.

## DATA AVAILABILITY

Matched raw ATAC-seq, H3K27ac ChIP-seq, RNA-seq, and WGS data from the A375, MM001, MM011, MM029, MM031, MM047, MM057, MM074, MM087, and MM099 patient-derived melanoma cell lines were obtained from European Genome-Phenome Archive (EGAS00001004136) and Gene Expression Omnibus (GSE134432, GSE142238, and GSE159965). Matched aligned ATAC-seq and WGS from TCGA GBM and LGG cohorts were downloaded from The National Cancer Institute (NCI) Genomic Data Commons (GDC) portal.

---

**Figure S1.**
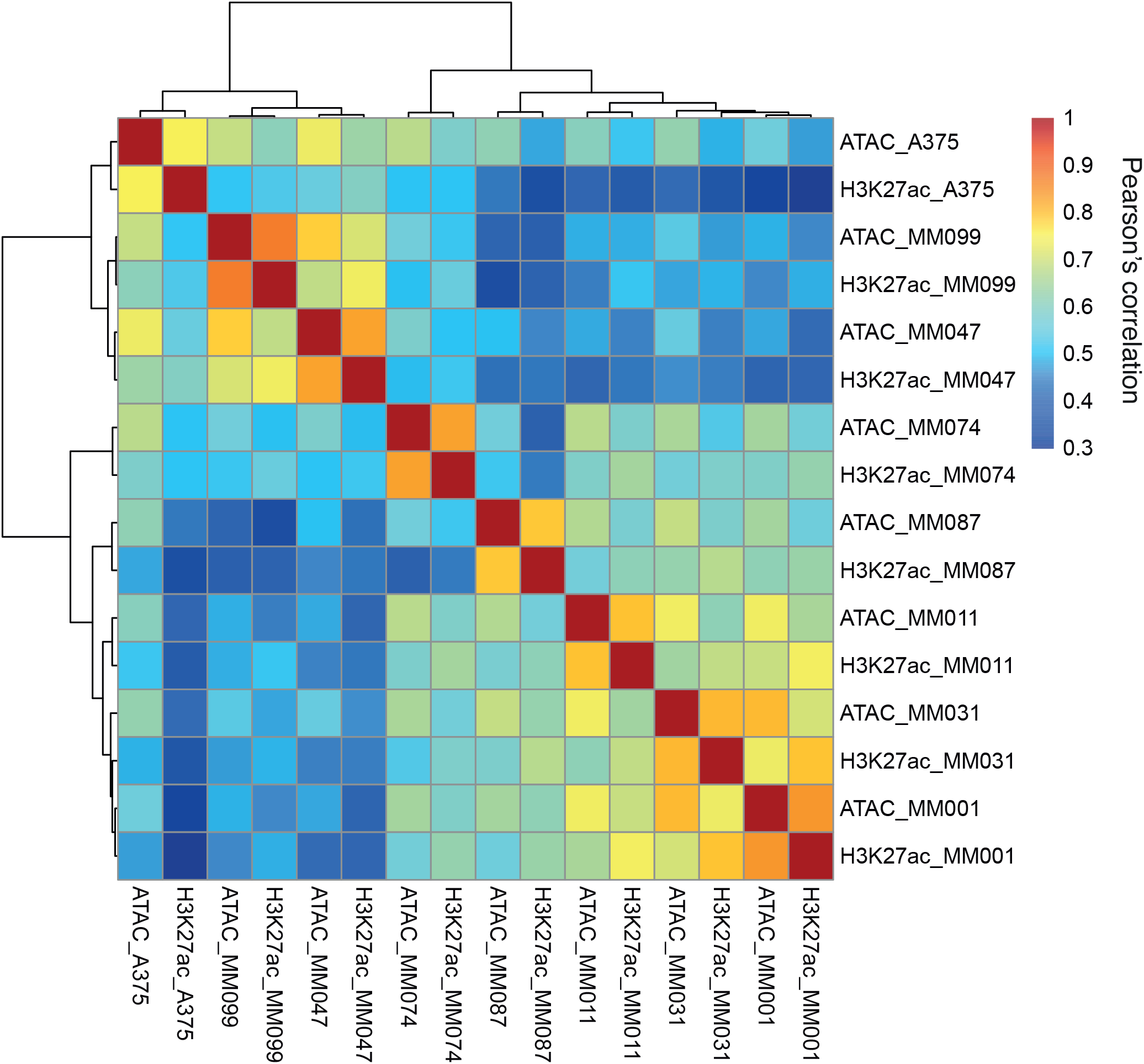
ATAC-seq links oncogenic alterations to coherent regulatory programs in patient-derived melanoma cell lines. Pearson correlation of signal intensity between ATAC-seq and H3K27ac ChIP-seq and 30× WGS, computed on the set of peaks identified with H3K27ac ChIP-seq. Rows and columns were hierarchically clustered using correlation distance and Ward’s D^2^ linkage.

**Figure S2.**
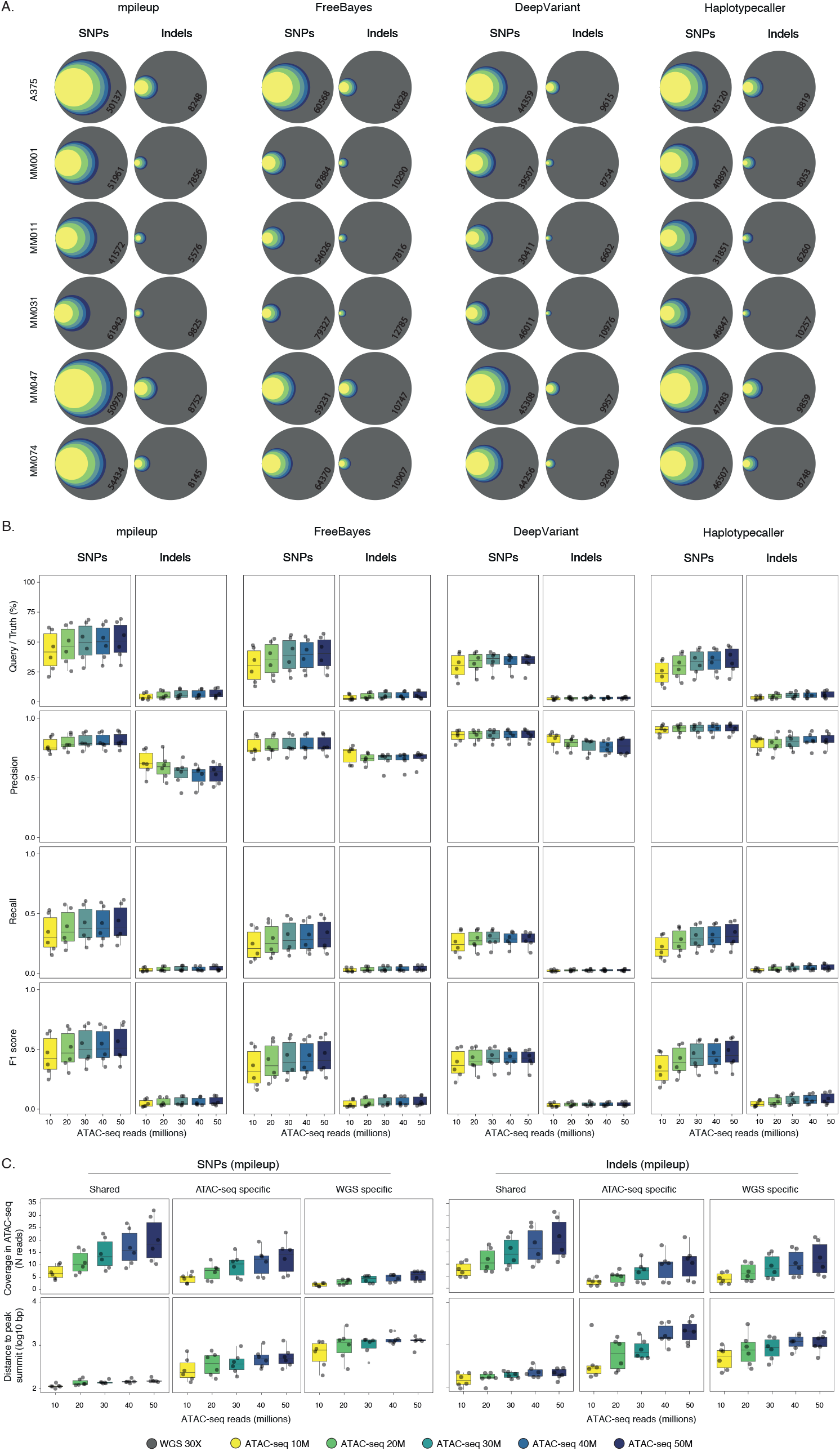
ATAC-seq delivers high-confidence SNPs within accessible chromatin in patient-derived melanoma cell lines. **(A)** Proportional Venn diagrams illustrating the overlap of SNPs and indels called within ATAC-seq peaks using bcftools/mpileup, FreeBayes, DeepVariant and GATK HaplotypeCaller, compared against matched 30× WGS, for each patient-derived melanoma cell line and ATAC-seq depth (10M-50M fragments). Concentric circles represent ATAC-seq callsets obtained at increasing sequencing depths (10M-50M fragments), and the outer grey circle represents the 30× WGS reference callset. Variants detected exclusively by only one technology across all depths were omitted to improve readability. The number displayed inside the grey circle corresponds to the total number of 30× WGS variants falling within the ATAC-seq accessible space, defined here as the peak set obtained with 50M ATAC-seq fragments. **(B)** (A) Boxplots illustrating the distribution of performance metrics for bcftools/mpileup, FreeBayes, DeepVariant and GATK HaplotypeCaller small-variant calls from ATAC-seq across patient-derived melanoma cell lines at each ATAC-seq sequencing depth (10M-50M fragments). Metrics include overlap with the matched 30× WGS callset, precision, recall and F1 score, computed using ATAC-seq calls as the query and 30× WGS as the truth set, and restricted to variants within ATAC-seq peaks. **(C)** Boxplots depicting the distribution of ATAC-seq coverage (top) and distance to peak summit (bottom) for shared variants, ATAC-seq-specific variants, and 30× WGS-specific variants, shown separately for SNPs (left) and indels (right), at each ATAC-seq sequencing depth (10M-50M fragments).

**Figure S3.**
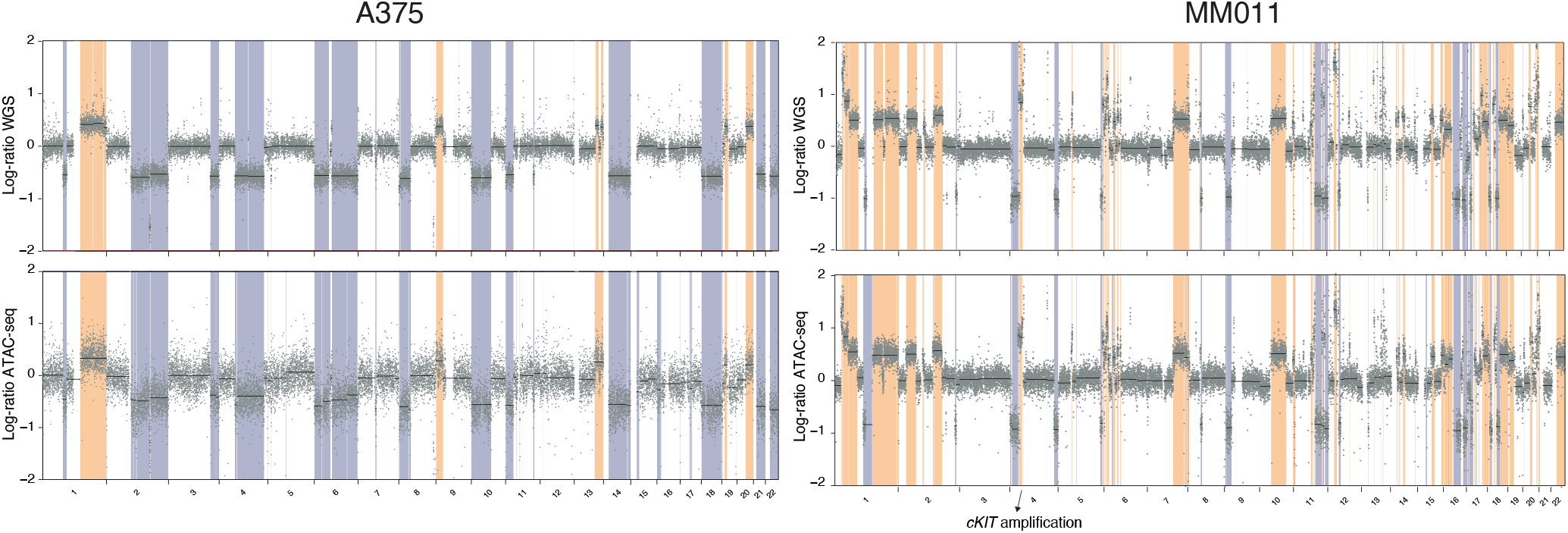
ATAC-seq recapitulates WGS copy-number alterations in patient-derived melanoma cell lines. Genome-wide segmented log_2_-transformed CNV profiles for representative patient-derived melanoma cell lines (A375 and MM011) inferred from 30× WGS (top) and ATAC-seq (bottom). Amplifications are highlighted in orange and deletions in purple. *cKIT* amplification in MM011 is indicated with an arrow.

**Figure S6.**
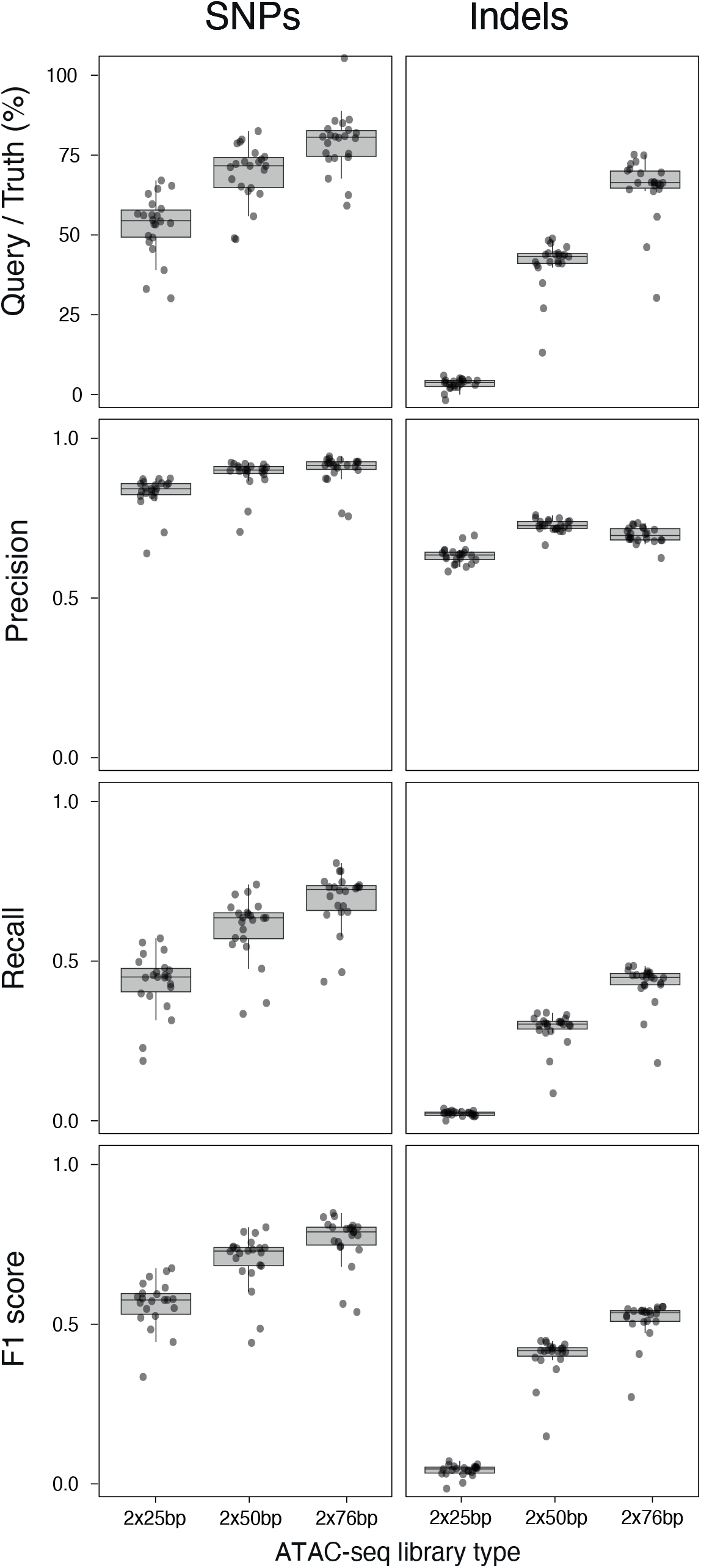
ATAC-seq delivers reliable small-variant calls within accessible regions and supports cohort genetic stratification in primary tumors. Boxplots illustrating the distribution of performance metrics for bcftools/mpileup small-variant calls from TCGA tumor ATAC-seq libraries, restricted to variants falling within ATAC-seq peaks. Libraries were downsampled to 50 million read pairs and reads were trimmed to 2×25 bp, 2×50 bp, or 2×75 bp prior to variant calling. Metrics (query/truth call ratio, precision, recall, and F1 score) were computed using ATAC-seq calls as the query set and the matched 30× WGS callset as the truth set.

**Table 1.** Dataset and sequencing metrics for the patient-derived melanoma cell line cohort and the TCGA LGG/GBM cohort.

**Table 2.** Small-variant counts and metrics by platform and caller for the patient-derived melanoma cell line cohort and the TCGA LGG/GBM cohort.

## Notes

### Competing Interest Statement

The authors have declared no competing interest.

### Summary of Updates

We have added a new analysis based on primary patient tumors from the TCGA low-grade glioma (LGG) and glioblastoma (GBM) cohort, profiled by bulk ATAC-seq with matched tumor whole-genome sequencing. This addition moves the study beyond patient-derived melanoma cell lines and anchors our conclusions in biologically and clinically heterogeneous human tumors. The revised manuscript now spans two complementary experimental settings. First, patient-derived melanoma cell lines provide a controlled context in which technical factors such as sequencing depth and read length can be explicitly examined. Second, primary LGG and GBM tumors introduce the complexity of real clinical material, including variable tumor purity, signal-to-noise ratios, and intratumor heterogeneity. The convergence of results across these two settings demonstrates that the genome-wide signals derived from ATAC-seq are not restricted to a favorable patient-derived cell line context but remain detectable and interpretable in patient samples. We have also refined the conceptual framing of the study. Our aim is not to present bulk ATAC-seq as a substitute for whole-genome sequencing, nor to claim comprehensive genome-wide variant detection from an accessibility-based assay. Instead, the revised manuscript delineates which classes of genomic information can be reliably extracted from bulk ATAC-seq, under which technical conditions, and with which inherent constraints. Throughout the results and discussion sections, we now more explicitly articulate the limits imposed by chromatin accessibility, read length, sparse allelic sampling, repetitive sequence content, and assay-specific biases. In addition, we have expanded the methodological descriptions, clarified the analytical pipelines - particularly for mitochondrial and telomere-associated analyses - and revised all figures to improve readability and interpretability. The discussion section has been strengthened to more critically address both the opportunities and the boundaries of genome-aware applications of bulk ATAC-seq, including implications for translational and clinical research. By showing that a single bulk ATAC-seq library can simultaneously inform regulatory landscapes, large-scale genome structure, accessible-region genotypes, mitochondrial allele-fraction profiles, and telomere-associated repeat abundance - across both cell lines and primary tumours - our work positions bulk ATAC-seq as a genome-aware, integrative backbone rather than a replacement for WGS. This perspective is particularly relevant in contexts where tissue availability, cost, or logistics limit the use of multiple genomic assays.

